# TDP-43 dysfunction induces cryptic circular RNAs in ALS/FTD

**DOI:** 10.64898/2026.09.17.752342

**Authors:** Dario Dattilo, Flaminia Pellegrini, Simone Barattucci, Anna-Leigh Brown, Jose Norberto S. Vargas, Ariana Gatt, Ryan Morrie, Georgiana Miller, Iris Bachmutsky, Zachary McEachin, Mingee Chung, Matthew J. Keuss, Eugeni Ryadnov, Matteo Zanovello, Puja R. Mehta, Francesca Mattedi, Michela Barioglio, Shubha Kamath, Sarah E Kargbo-Hill, Joanna Palade, Isabelle Kowal, Jonathan Glass, Marla Gearing, Edward B Lee, Melissa E Murray, Dennis W Dickson, NYGC ALS Consortium, Eric M Green, Nicholas T Seyfried, Sanjay Chandriani, Michael Ward, Pietro Fratta

## Abstract

Nuclear depletion of TDP-43 is a defining pathological feature of amyotrophic lateral sclerosis (ALS) and frontotemporal dementia (FTD), leading to widespread RNA misprocessing, including the formation of cryptic exons. Here, we identified TDP-43 as a regulator of circular RNA (circRNA) biogenesis in multiple human neuronal cell models, and showed that its dysfunction induces the de novo formation of cryptic circular RNAs (c-circRNAs). Analysis of post-mortem brain transcriptomic data identified a subset of c-circRNAs which are specific for ALS and FTD cases with TDP-43 pathology. Further, we developed highly sensitive rolling-circle amplification-based circRNA detection assays that allow to distinguish TDP-43 pathology in human CNS tissues with a 0.99 AUC. We found that c-circRNAs can co-occur with cryptic linear splicing events, uncovering complex RNA misprocessing hotspots that induce loss of disease-relevant proteins, including *RPTOR* and *EHMT1*. Notably, one of these c-circRNAs originates from *UNC13A*, a gene whose cryptic exon has previously been linked to one of the major GWAS hits in ALS/FTD and that is being pursued as a therapeutic target through splice-switching ASOs. We showed that c-circUNC13A is co-regulated with the linear cryptic transcript and suppression of UNC13A cryptic exon results in c-circUNC13A reduction in cultured neurons and in vivo, highlighting its potential as a target engagement biomarker for emerging UNC13A-directed therapies. Overall, this work identifies a novel molecular mechanism for TDP-43 dysfunction, opening novel avenues for understanding disease pathogenesis and developing much needed pathology biomarkers.

## Introduction

Cytoplasmic accumulation and nuclear depletion of TDP-43 are the crucial hallmark of amyotrophic lateral sclerosis (ALS) and approximately 50% of frontotemporal dementia (FTD) cases^1,2^. TDP-43 is an RNA binding protein involved in multiple aspects of RNA processing, including splicing and polyadenylation^3^. Nuclear TDP-43 loss leads to the formation of novel splicing and termination events (cryptic events; CEs)^4–7^. The majority of CEs lead to protein loss either through nonsense-mediated-decay (NMD) or transcript truncation, and in some cases cryptic polyadenylation induces protein upregulation through increased RNA stability^4–6^. In a minority of cases, CEs are in frame and do not contain premature stop codons, giving rise to additional peptides named cryptic peptides (CP)^8,9^.

Cryptic RNA processing alterations have been shown to be relevant to disease pathogenesis and progression^10,11^, and targeted therapeutics aiming to rescue single CEs are currently in clinical trials^12–14^. CEs and CPs can be exceptionally specific to the disease condition and have been used to detect and quantify the extent of TDP-43 dysfunction in patient tissue^15,16^. This specificity also has potential for biofluid biomarker development. However CEs are unstable in biofluids and are often degraded intracellularly^15,17^. CPs have been detected in CSF and plasma^8,9^, but their typical low abundance may limit detection through mass-spectrometry and antibody-based assays, highlighting the need for more specific and sensitive assays. As a result, robust biomarkers of TDP-43 dysfunction remain a critical gap in the field.

Circular RNAs (circRNAs) are a class of endogenous RNAs characterized by a covalent closed loop structure^18^. Such molecules have gained attention over the last 10 years thanks to the advancements in sequencing preparation methods and analysis^18^, which have revealed their ubiquitous expression across eukaryotes and their enrichment in neuronal tissues^19^. Circular RNA levels have also been reported to be dysregulated in many pathological conditions, including Alzheimer’s disease^20,21^. Whether TDP-43 dysfunction can impact on circRNA expression has not yet been addressed.

We found that loss of TDP-43 in human cultured neurons leads to widespread changes in circRNAs and also to the formation of novel circRNAs (cryptic circRNAs, c-circRNAs). These c-circRNAs accumulate selectively in cortical and spinal cord regions affected by TDP-43 pathology in FTD and ALS, discriminating TDP-43 proteinopathies with very high sensitivity and specificity. We found that c-circRNAs can co-occur with other CEs within the same genes, but are more stable and readily detectable than their linear counterparts. These complex RNA misprocessing events can also result in the loss of proteins with key roles in biological pathways disrupted in neurodegeneration, such as autophagy, as exemplified by RPTOR^22,23^. Lastly, we identify a c-circRNA in the *UNC13A* locus, where a cryptic exon associated with ALS disease progression^24,25^ is also detected and is currently being targeted in clinical trials^13,14^. Our work shows that the UNC13A c-circRNA and cryptic exon are co-regulated and that therapeutic ASOs correcting the cryptic exon also rescue the c-circRNA, supporting its potential as target engagement biomarker for splice site-switching approaches.

## Results

### TDP-43 loss induces differential circRNA expression

TDP-43 takes part in multiple RNA processing steps, and we asked whether it also regulates circRNA levels and biogenesis. We investigated total RNA-seq from four neuronal TDP-43 knockdown experiments: two carried out in neuroblastoma cell lines (SH-SY5Y and SK-N-BE(2)) and two in hiPSC-derived neuronal models, namely lower motor neurons (i3LMN) and cortical neurons (i3N) (**Fig. S1A and S1B**). Using the CIRI2 tool^26^ we identified numerous circRNAs: 4761 in SH-SY5Y, 2204 in SK-N-BE(2), 6939 in i3LMN, and 10097 in i3N (**Fig. S1C**) with 1552 detected across all 4 datasets (**Fig. S1D**). In short-read RNA sequencing, circRNA quantification relies on the detection of the back-splicing junction (BSJ), the only region being unequivocally attributed to a circRNA, therefore significantly reducing the sensitivity of circRNA detection compared to linear RNA counterparts in conventional RNA-seq. As a result, the overlap of circular species detected across datasets is lower than the linear ones (**Fig. S1D**). Nonetheless, differential expression analysis revealed significant circRNAs changes (**Fig. 1A**), with a total of 1118 distinct circRNAs upregulated (**Fig. 1B**) and 1152 downregulated (**Fig. S1E**) across the four datasets.

**Figure 1.**
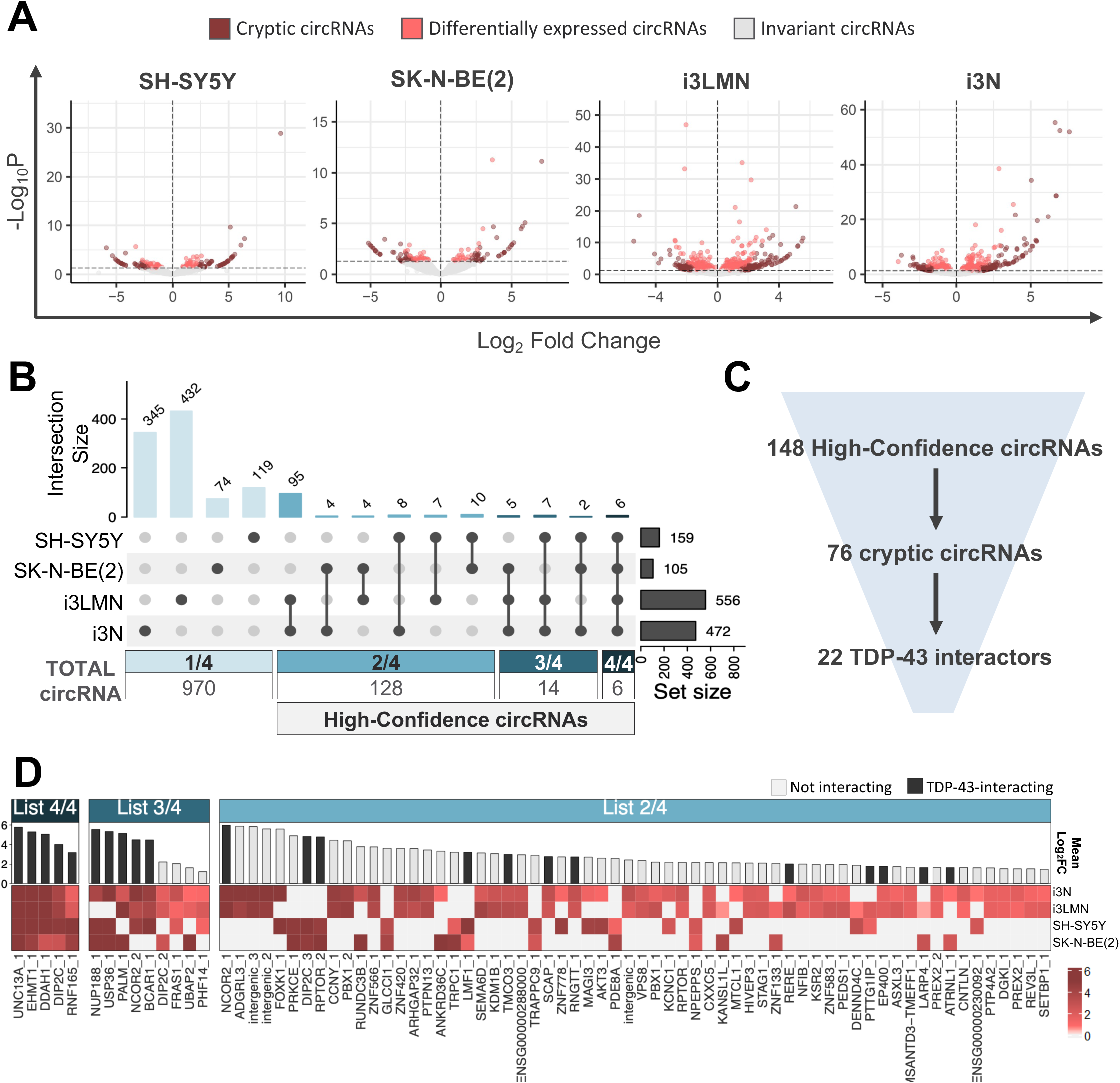
TDP-43 depletion drives c-circRNAs expression in multiple cell models: **A)** Volcano plots showing the log₂ fold change (x-axis) and −log₁₀ P value (y-axis) for all circRNAs identified in the four RNA-seq datasets (SH-SY5Y, SK-N-BE(2), i3LMNs and i3Ns) following TDP-43 knockdown. Invariant circRNAs are shown in grey, differentially expressed circRNAs (points above the horizontal −log10 P value threshold) in red, and cryptic circRNAs in dark red. **B)** UpSet plot showing the overlap of upregulated circRNAs identified across the four TDP-43 knockdown models: SH-SY5Y, SK-N-BE(2), i3LMN, and i3N. Horizontal bars on the right indicate the total number of circRNAs upregulated upon TDP-43 knockdown in each model, whereas vertical bars represent the size of each intersection. Connected dots denote the datasets contributing to each intersection. The lower panel summarises the number of upregulated circRNAs according to the number of models in which they were identified. CircRNAs upregulated in at least two of the four models were classified as “High-Confidence circRNAs” and retained for downstream analyses. **C)** Funnel diagram illustrating the filtering strategy used to identify TDP-43-associated cryptic circRNAs. Among the 148 High-Confidence circRNAs, 76 fulfilled the criteria for cryptic circRNAs (c-circRNAs). Integration with TDP-43 CLIP datasets identified 22 c-circRNAs with evidence of TDP-43 binding within their flanking introns. **D)** Heatmap showing the Log_2_FoldChange (Log_2_FC) values of High-Confidence c-circRNAs across the four TDP-43 knockdown cellular models: SH-SY5Y, SK-N-BE(2), i3LMNs, and i3N. c-circRNAs are grouped according to the number of datasets in which they were identified (List 2/4, List 3/4, and List 4/4) and ranked by their mean Log₂FC across datasets, as indicated by the bar plots above the heatmap. Colour intensity indicates the magnitude of circRNA upregulation following TDP-43 depletion. Black bars denote c-circRNAs with evidence of TDP-43 binding within their flanking introns based on CLIP data, whereas grey bars indicate the absence of detectable TDP-43 binding.

To understand whether changes are driven by alterations in pre-mRNA levels or are directly induced by changes in back-splicing, we examined the expression of the corresponding linear isoforms and classified circRNAs as “invariant”, if the corresponding linear isoform showed no significant variation, “concordant”, if both varied in the same direction, and “discordant” if they varied in opposite directions. Across all lines, the majority of circRNAs (65–97%) fell into the “invariant” or “discordant” categories, indicating that circRNA changes are not determined by alterations in transcription (**Fig. S2A**). Analysis of changes in the circular-to-linear ratio (ΔCLR) between TDP-43 knockdown and control conditions showed that changes in circRNA abundance closely mirrored changes in ΔCLR, further reinforcing that differential circRNA expression primarily reflects altered back-splicing efficiency rather than changes in host gene expression (**Fig. S2B**).

We assessed whether circRNA upregulation was shared across the different cell types and found that out of the 1118 found to be changed, 970 were specific to a single cell line, while 148 were upregulated in at least two of the four models (**Fig. 1B**). The low degree of overlap was primarily due to their lack of detection in the other cellular models (**Fig. S3A and S3B**). Throughout the manuscript, we refer to circRNAs changed in at least 2 datasets as “High-Confidence circRNAs” (**Fig. 1C**).

We next asked whether TDP-43 regulates back-splicing through direct RNA binding and investigated the presence of TDP-43 binding sites in the introns flanking the back-splicing exons, as these are known to play a critical role in circularization^27,28^ (**Fig. S3C**). Using publicly available TDP-43 CLIP-seq data^29^ (**Fig. S3D**), we observed a significant enrichment of TDP-43 binding in at least one of the two flanking introns of upregulated circRNAs compared with a control set of circRNAs that do not change after TDP-43 knockdown (**Fig. S3E and S3F**). Notably, the enrichment was more prominent for those circRNA species being upregulated across multiple cellular models (**Fig. S3F**).

### TDP-43 loss promotes cryptic circRNA (c-circRNA) formation

Amongst the upregulated circRNAs, we identified a subset detected exclusively following TDP-43 knockdown and absent in control conditions. This reveals a previously unrecognised consequence of TDP-43 dysfunction: the de novo emergence of circular RNAs. As this mirrors the definition of cryptic events^7^, we classified this class as “cryptic circRNAs” (c-circRNAs) (**Fig. 1A**). In total, we identified 543 c-circRNAs (**Fig. S3G**), 76 of which belong to the High-Confidence group (**Fig. 1C**). These High-Confidence cryptic circRNAs are shown in Fig. 1D, ranked according to the number of lines in which each circRNA was identified and their magnitude of induction upon TDP-43 knockdown. Notably, 22 of these circRNAs exhibited direct TDP-43 binding (**Fig. 1C and 1D**).

### c-circRNAs specifically associate with TDP-43 pathology in ALS and FTD post-mortem CNS

The detection of differentially-expressed transcripts due to TDP-43 dysfunction in bulk postmortem tissue can be challenging, as only a minority of cells exhibit TDP-43 nuclear loss. On the other hand, detection of cryptic events is a more reliable measure, as these are absent in physiological conditions. We therefore investigated the presence of c-circRNAs in New York Genome Center (NYGC) ALS Consortium RNA-seq data. This cohort includes different central nervous system (CNS) regions from postmortem tissues of ALS and FTD cases, along with non-disease controls, overall comprising 1,863 samples (**Fig. S4A**). To minimize the impact of confounding factors, we verified that pathological samples did not differ significantly from controls in terms of library depth, RNA integrity number (RIN), age at death, or sex (**Fig. S4B–E**).

We used CIRI2 to quantify circRNAs expressed in control and ALS-TDP samples, and detected over half a million circRNAs (**Fig S5A and S5B**). We performed an enrichment analysis to identify circRNAs detected in a higher number of ALS-TDP samples compared to controls (**Fig. S5C and S5D**). Within this group, we defined *c-circRNAs* as those expressed in less than 5% of control samples (**Fig. S5C**). Overall, we identified 466 different c-circRNAs (**Fig. S5E**) and, among these, 65 displayed at least three fold enrichment in ALS-TDP versus control samples (**Fig 2A and S5F**).

**Figure 2.**
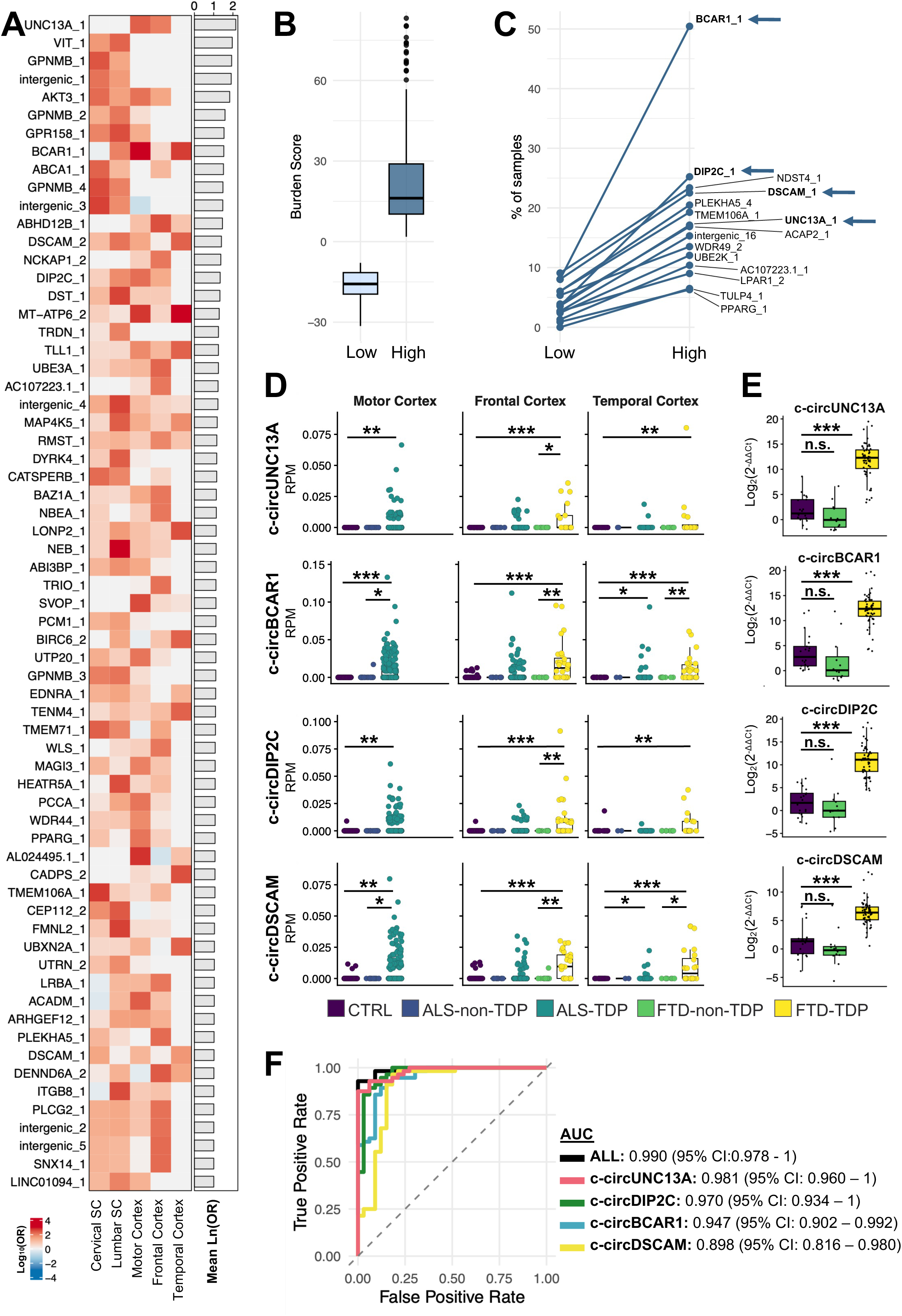
c-circRNAs sensitively discriminate TDP-43 proteinopathy in postmortem ALS/FTD: **A)** Heatmap showing the enrichment (natural logarithm of the odds rat, LnOR) of cryptic circRNAs (c-circRNAs) in ALS-TDP samples relative to controls across the five CNS regions analysed (cervical spinal cord, lumbar spinal cord, motor cortex, frontal cortex and temporal cortex). c-circRNAs are ranked by their median LnOR across tissues, as indicated by the bar plots to the right of the heatmap. Colour intensity reflects the magnitude of c-circRNA enrichment, calculated using Fisher’s exact test. **B)** Box plot showing the distribution of TDP-43 cryptic burden score in ALS-TDP samples assigned to the low- and high-burden groups. Samples were stratified according to the lowest and highest quartiles of the TDP-43 cryptic burden score, respectively. **C)** Line plot showing the proportion of ALS-TDP samples expressing each c-circRNA in the low- and high-burden groups. Each line represents one c-circRNA. c-circRNAs significantly enriched in the high-burden group (*P* < 0.05, Fisher’s exact test) are labelled. No c-circRNAs were significantly enriched in the low-burden group. Blue arrows indicate the four candidate c-circRNAs selected for further validation. **D)** Box plots showing the expression of selected c-circRNAs (c-circUNC13A, c-circBCAR1, c-circDIP2C and c-circDSCAM) in the NYGC ALS Consortium RNA-seq dataset across disease groups and CNS regions. Expression of the c-circRNAs was quantified from back-splice junction (BSJ) counts normalised for library size (reads per million, RPM). Each dot represents an individual sample. Statistical significance was assessed using Wilcoxon-Mann-Whitney test with Bonferroni correction (*p < 0.05, **p < 0.01, ***p < 0.001). Non-significant comparisons (p > 0.05) are not reported. **E)** RT–qPCR analysis of selected c-circRNAs (c-circUNC13A, c-circBCAR1, c-circDIP2C, c-circDSCAM) in post-mortem human frontal cortex samples from healthy controls (CTRL, n = 21) and FTD patients with (FTD-TDP, n = 56) or without (FTD-non-TDP, n = 12) TDP-43 pathology. c-circRNA expression was normalised to GAPDH mRNA and is presented as fold change (log₂(2^−ΔΔCt)), with respect to a control sample set to a value of 1. Each dot represents an individual sample. Statistical significance was assessed using Wilcoxon-Mann-Whitney test with Bonferroni correction (n.s. p > 0.05, ***p < 0.001). **F)** Receiver operating characteristic (ROC) curves evaluating the ability of individual c-circRNAs and a combined four-marker model to discriminate FTD-TDP from healthy controls and FTD cases without TDP-43 pathology. Areas under the curve (AUCs) are indicated.

We then tested whether c-circRNA abundance was responsive to the degree of TDP-43 dysfunction, and used the TDP-43 cryptic burden score^15^, to estimate the extent of TDP-43 nuclear loss-of-function. We classified samples into a “low burden” group (lowest burden score quartile) and a “high burden” group (highest burden score quartile) (**Fig. 2B and S6A**), ensuring equal sample size between groups and confirming the absence of differences in sequencing depth (**Fig. S6B**). Among the identified c-circRNAs, none were enriched in the low burden group, whilst 15 were significantly enriched in the high burden group (**Fig. 2C**), identifying them as good markers of TDP-43 pathology.

Amongst the 15 TDP-43 responsive c-circRNAs, *c-circUNC13A*, *c-circBCAR1*, and *c-circDIP2C* were identified in all our cellular models (**Fig. 2C**) and are bound by TDP-43 within their flanking introns (**Fig. S7A**). Another relevant candidate was *c-circDSCAM*, which displays a cryptic expression profile in five out of five tissues analysed (**Fig. 2A**) and is among the most enriched circRNAs in the high burden group (**Fig. 2C**). We confirmed that all four c-circRNAs show specific enrichment in both ALS and FTD samples with TDP-43 pathology (ALS-TDP and FTD-TDP), when compared not only with healthy controls but also with individuals affected by non–TDP-43 forms of ALS (associated with *SOD1* or *FUS* mutations) or FTD (associated with TAU or FUS) (**Fig. 2D and S7B**). Importantly, none of these c-circRNAs are associated with a significant increase in the expression of their corresponding linear isoforms (**Fig. S7C**). Based on their robust detection across models and disease specificity, we selected these four c-circRNAs for targeted validation in post-mortem tissue.

### c-circRNAs enable sensitive detection of TDP-43 pathology in tissue

A major challenge when analysing post-mortem brain tissue is that cells harbouring TDP-43 pathology represent only a small minority. Indeed, even highly abundant cryptic events such as the STMN2 cryptic exon are often undetected or supported by only a few reads in bulk RNA-seq datasets^30^. To overcome this limitation, we developed an amplification strategy and targeted assays to enhance c-circRNA detection in bulk tissue. We used the Group II intron reverse transcriptase (RT) Induro, which performs efficient rolling-circle reverse transcription (RC-RT)^31^ (**Fig. S8A**) generating long cDNA molecules with multiple copies of the BSJ, therefore enhancing circRNA detection (**Fig. S8B**). Direct comparison of Induro with SuperScript IV, a reverse transcriptase unable to perform rolling-circle amplification, showed comparable qPCR signals for a linear RNA template, whereas Induro increased the signal from a circular RNA template by more than 20-fold (**Fig. S8B and S8C**), demonstrating that RC-RT selectively enhances circRNA, but not linear RNA, detection.

We applied RC-RT followed by BSJ-specific TaqMan assays on frontal cortex RNAs from healthy controls (n=21) and FTD-TDP patients with (GRN n=17, C9-FTD n=14, sporadic FTLD-TDP n=25) or without TDP-43 pathology (FTD-FUS n=12). All four selected c-circRNAs displayed a markedly elevated signal in FTD-TDP samples compared to FTD-FUS and healthy controls (**Fig. 2E**). Importantly, this difference was detected in all FTD-TDP subtypes (**Fig. S8E**).

To assess the ability of these c-circRNAs to distinguish FTD-TDP from controls and FTD-non-TDP cases, we performed ROC analyses. Each of the four c-circRNAs demonstrated strong discriminatory power (AUC 0.90–0.98), and the combined model achieved an AUC of 0.99, indicating high predictive accuracy and supporting their potential as biomarkers of TDP-43 proteinopathy (**Fig. 2F**).

### c-circRNAs reveal complex misprocessing events linked to neurodegeneration

The c-circRNA most predictive of TDP-43 pathology occurs in *UNC13A*, a gene where TDP-43 loss also induces a linear CE that is associated with one of the strongest GWAS signals in both ALS and FTD^24,25,32–34^. This event consists in the aberrant inclusion of a cryptic exon within intron 20, corresponding to one of the flanking introns of the aforementioned c-circUNC13A. This event consists in the aberrant inclusion of a cryptic exon within intron 20, corresponding to one of the flanking introns of the aforementioned c-circUNC13A. We therefore asked whether any other linear CE occurred within the flanking introns of the TDP-43-induced c-circRNAs (**Fig. S9A**). As CEs often trigger NMD, which can mask their detection^15,35^, we utilised a recently compiled comprehensive list of CEs generated from multiple datasets, including the ones where NMD was inhibited^15^. We found that 14 out of the 76 High-Confidence c-circRNAs identified in cell lines harbored CE within one of their flanking introns (**Fig. 3A**). These cases of aberrant linear splicing included both exon inclusion and skipping events.

**Figure 3.**
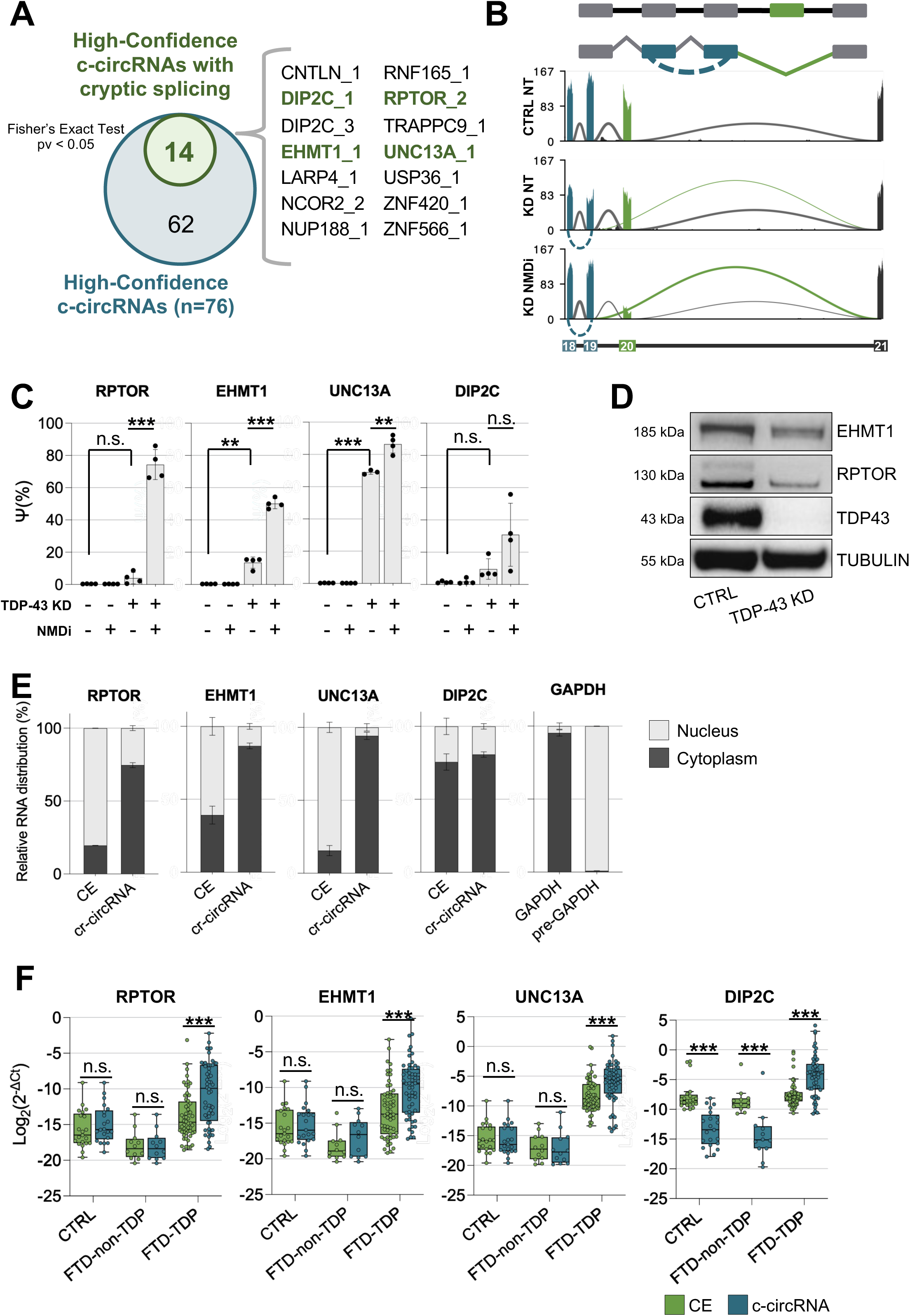
c-circRNAs and linear CEs associate in genes linked to neurodegeneration: **A)** Venn diagram showing the overlap between the High-Confidence cryptic circRNAs (blue) and the High-Confidence cryptic circRNAs with a cryptic splicing event in one of their flanking introns (green). Fisher’s exact test was used to assess the significance of the overlap. The 14 common circRNAs are listed on the right. c-circRNAs shown in bold were selected for further analyses. **B)** Sashimi plot showing the concomitant occurrence of a cryptic splicing event (green) and a cryptic circular RNA (c-circRNA)–forming event (blue) in the *RPTOR* gene. The upper panel shows a schematic representation of the events, while the lower panels display read coverage and splice junction usage under the indicated conditions: control (CTRL NT), TDP-43 knockdown (KD NT), and TDP-43 knockdown with nonsense-mediated decay inhibition (KD NMDi). **C)** Percentage spliced in (PSI, ψ) values for the linear cryptic splice isoforms of UNC13A, RPTOR, EHMT1, and DIP2C relative to their corresponding correctly spliced junctions. ψ values were calculated from bulk RNA-seq data generated from SH-SY5Y cells under control or TDP-43 knockdown conditions, with or without cycloheximide-mediated inhibition of nonsense-mediated decay (NMD). Bars represent the mean ± SD of independent biological replicates, with individual replicates shown as black dots (n=4). Statistical significance was assessed using a two-tailed unpaired Student’s *t* test with correction for multiple test comparison (FDR Benjamini–Hochberg) (n.s. p > 0.05, **p < 0.01, ***p < 0.001). **D)** Representative western blot showing EHMT1, RPTOR, and TDP-43 protein levels in control and TDP-knockdown i3N; α-Tubulin (TUBULIN) was used as loading control. n = 3 independent biological replicates. **E)** Subcellular localisation of linear cryptic transcripts (CE) transcripts and the corresponding cryptic circRNAs (c-circRNA) derived from the indicated genes. Nuclear and cytoplasmic fractions were isolated from TDP-43-depleted SH-SY5Y cells, and transcript abundance was quantified by RT–qPCR. Stacked bar plots show the proportion of each transcript detected in the nuclear (light grey) and cytoplasmic (dark grey) fractions. GAPDH and pre-GAPDH were used as cytoplasmic and nuclear controls, respectively. Bars represent the mean ± SD of three independent biological replicates (n = 3). **F)** RT–qPCR analysis linear cryptic transcripts (CE, green) and their corresponding cryptic circRNAs (c-circRNA, blue) in post-mortem human frontal cortex samples from healthy controls (CTRL, n = 21) and FTD patients with (FTD-TDP, n = 56) or without (FTD-non-TDP, n = 12) TDP-43 pathology. Transcript abundance was normalised to GAPDH mRNA and is presented as log₂-transformed expression values (log₂(2^−ΔCt)). Each dot represents an individual sample. Statistical significance was assessed using Wilcoxon-Mann-Whitney test with Bonferroni correction (n.s. p > 0.05, ***p < 0.001).

Intriguingly, within this subset we identified not only *UNC13A* (**Fig. S9B**) but also two other genes linked to neurological diseases: *RPTOR* (**Fig. 3B**) and *EHMT1* (**Fig. S9C**). *RPTOR* encodes a crucial scaffolding protein for the formation and regulation of the mTORC1 signalling pathway, a central regulator in neuronal homeostasis which has been associated with the progression of neurodegenerative diseases^22,23^, and *EHMT1* encodes a histone methyltransferase that plays a critical role in brain development and cognitive function, with mutations giving rise to Kleefstra syndrome^36,37^.

*UNC13A*, *RPTOR* and *EHMT1* transcripts are all sensitive to NMD and are downregulated upon TDP-43 depletion and rescued upon NMD inhibition (**Fig. S10A**). Whilst the *UNC13A* CE can be reliably observed in both cell culture experiments and post-mortem bulk RNA, the *RPTOR* and *EHMT1* CEs are extremely vulnerable to (**Fig. 3C**) and have therefore been defined as “dark” (i.e., only identifiable following NMD inhibition in vitro) and not detected in NYGC post-mortem tissues^15^. We validated the NMD sensitivity of *UNC13A*, *RPTOR*, and *EHMT1* cryptic isoforms by RT-qPCR in SH-SY5Y cells following TDP-43 depletion, while demonstrating that the corresponding c-circRNAs are resistant to NMD (**Fig. S10B and S10C**).

Whilst the impact of *UNC13A* mis-splicing has been characterised elsewhere^14,24,25^, we sought to assess the impact of the *RPTOR* and *EHMT1* multi-misprocessing events and investigated whether these result in reduced protein expression upon TDP-43 depletion. Western blot on lysates from iPSC-derived cortical neurons showed a strong down-regulation of RPTOR and EHMT1 proteins as an effect of TDP-43 depletion (**Fig. 3D and S3D**).

Overall these results highlight how c-circRNAs, which are insensitive to NMD, can unveil more complex mis-splicing events which have significant consequences on disease-related protein expression.

### c-circRNAs have a distinct subcellular localization and are better detected compared to their CE counterparts

As c-circRNAs and CEs are distinct transcripts deriving from the same gene, we decided to compare their subcellular localisation. We therefore investigated the distribution of the disease-relevant c-circRNAs and their CE counterparts by performing nucleo-cytoplasmic fractionation in TDP-43 depleted SH-SY5Y cells. All CEs containing a premature stop codon were predominantly localised in the nuclear fraction, whilst all c-circRNAs were predominantly localised to the cytosolic fraction (**Fig. 3E**). This result supports the potential of c-circRNAs to be incorporated within extracellular vesicles.

We then compared the detectability of c-circRNAs and their CE counterparts in post-mortem tissues. We performed rolling-circle RT followed by RT-qPCR analysis and found an increase of all circular and linear targets in FTD-TDP compared to controls and non-TDP-FTD samples - therefore confirming their cryptic nature - with the exception of *DIP2C* CE that remained unchanged (**Fig. 3F**).

Notably, while c-circRNAs and their corresponding CEs were detected at comparable levels in controls and non-TDP FTD samples, c-circRNAs displayed consistently enhanced detectability in FTD-TDP tissues (**Fig. 3F**). To quantify this effect, we calculated a c-circRNA:CE ratio for each sample. The ratio favoured the circRNAs exclusively in FTD-TDP samples, confirming the reliability of the signal rather than a general amplification effect introduced by the assay (**Fig. S10E**).

These results corroborate that c-circRNA molecules show superior potential over their linear counterparts as biomarkers in clinical samples, likely due to their higher resistance to exonucleases, insensitivity to NMD, cytoplasmic localization and the technical advantage given by rolling circle amplification.

### c-circUNC13A is more stable than UNC13A cryptic exon

Among the c-circRNAs, c-circUNC13A is of particular interest because of its proximity to the UNC13A CE and the GWAS risk SNPs^24,25,32–34^, and we therefore analysed its features in depth. While UNC13A produces several circular isoforms, with some of them increasing upon TDP-43 depletion (**Fig. S11A**), only the isoform containing exons 21 to 30 (previously referred to as “*c-circUNC13A*”) fulfilled criteria of being cryptic across all analysed datasets (**Fig. 4A and Fig. S11B**). We validated this by RT-qPCR across the four cell lines (**Fig. 4B and Fig. S11C**), and also its circular nature by showing it to be resistant to RNase R treatment, as opposed to linear UNC13A CE (**Fig. S11D**).

**Figure 4.**
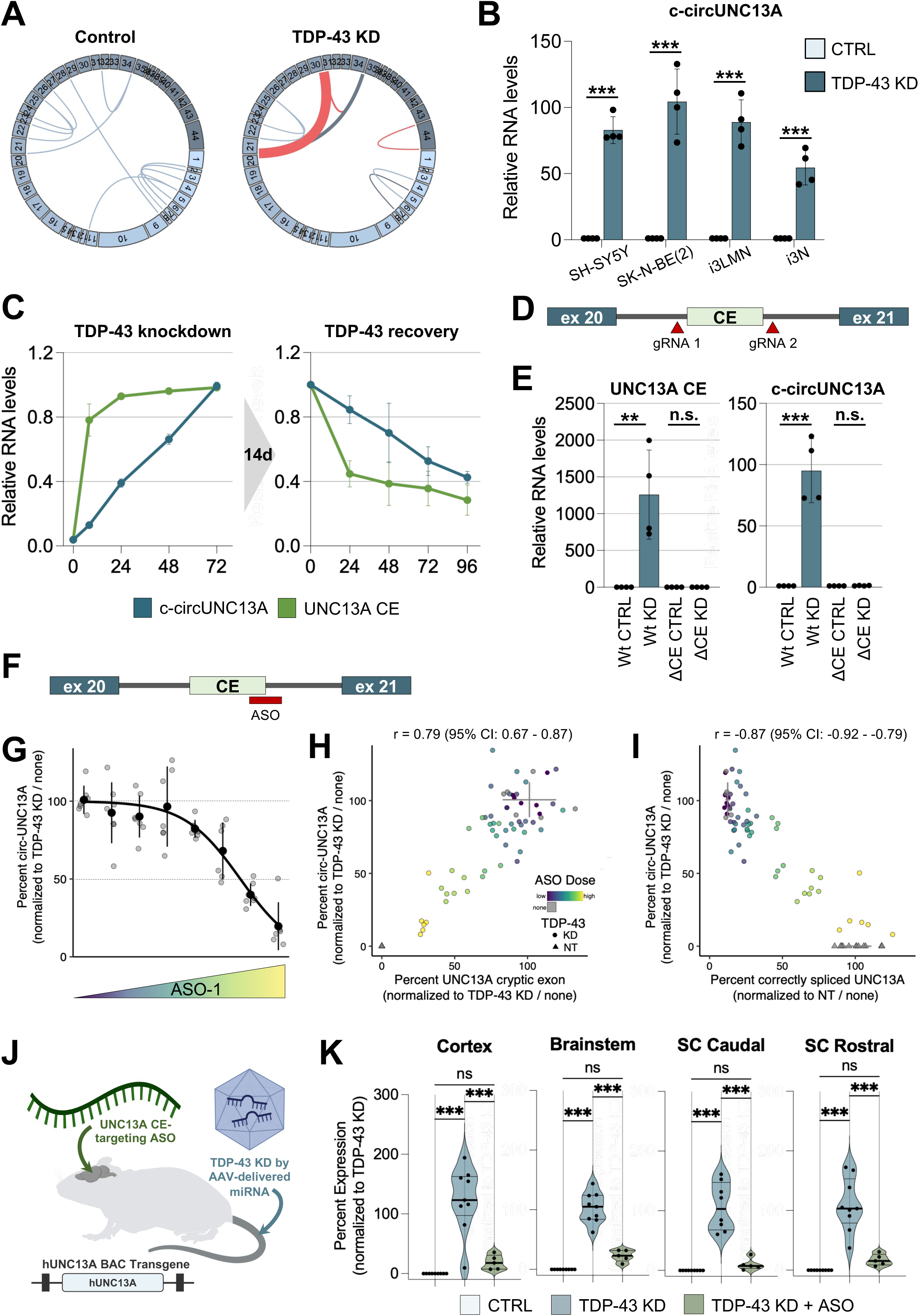
c-circUNC13A is co-regulated with UNC13A CE in vitro and in vivo: **A)** Schematic representation of UNC13A circRNA isoforms detected across the four cellular models under control (Control) and TDP-43 depletion (TDP-43 KD). Blue arcs indicate circRNA isoforms detected under both conditions, whereas red arcs indicate circRNA isoforms detected exclusively following TDP-43 depletion. Arc thickness is proportional to the mean circRNA expression level (CPM) across the four datasets. **B)** RT–qPCR analysis of c-circUNC13A expression across the four cellular models indicated under control (CTRL) and TDP-43 knockdown (TDP-43 KD) conditions. Transcript abundance was normalised to GAPDH mRNA and is presented as relative expression values (2^−ΔΔCt), with the control condition set to a value of 1. Each dot represents an independent biological replicate (n=4). Statistical significance was assessed using a two-tailed unpaired Student’s *t* test (***p < 0.001). **C)** RT–qPCR analysis of UNC13A cryptic exon (CE) and c-circUNC13A abundance during PROTAC-mediated TDP-43 depletion and subsequent recovery. Left panel: transcript abundance was measured at the indicated time points following induction of TDP-43 depletion using PROTAC. Right panel: after 14 days of PROTAC treatment, cells were switched to the catalytically inactive competitor (PROTACi) to restore TDP-43 levels, and transcript abundance was measured at the indicated recovery time points. In both panels, transcript abundance was normalised to GAPDH mRNA and is presented as relative expression values (2^−ΔΔCt), with the final time point of PROTAC treatment (72 h) and the initial time point of PROTACi treatment (0 h) each set to a value of 1. Data represent the mean ± SD of three independent biological replicates (n=3). **D)** Schematic representation of the CRISPR/Cas9-mediated deletion of the UNC13A cryptic exon (CE, green). The positions of the two guide RNAs (gRNA1 and gRNA2) used to generate the ΔCE cell line are indicated. **E)** RT–PqCR analysis of the indicated targets in SH-SY5Y cells with a wild-type (Wt) or UNC13A cryptic-exon deleted (ΔCE) background, under control (CTRL) or TDP-43 knockdown (KD) conditions. Transcript abundance was normalised to GAPDH mRNA and is presented as relative expression values (2^−ΔΔCt), with respect to the CTRL samples set to a value of 1. Each dot represents an individual sample (n=4). Statistical significance was assessed using a two-tailed unpaired Student’s *t* test (n.s. p > 0.05, **p < 0.01,***p < 0.001). **F)** representation of a high-performing splice-correcting antisense oligonucleotide (ASO) targeting the UNC13A cryptic exon (CE, green). The ASO binding site is indicated in red. **G)** Dose–response curve showing the effect of increasing concentrations of ASO-1 on c-circUNC13A expression in TDP-43-depleted cells. c-circUNC13A levels are presented relative to the TDP-43 knockdown condition, set to 100%. Individual data points are shown with mean ± SD. Data is fit with a 4 parameter log-logisitic function. **H)** Correlation between ASO-1-mediated reductions in c-circUNC13A and UNC13A cryptic exon expression. Data points are coloured according to the ASO-1 concentration. Crosshairs indicate the c-circUNC13A and UNC13A cryptic exon levels in TDP-43 KD only cells (mean ± SD for each axis). Triangles indicate basal conditions in the absence of TDP-43 knockdown, where neither c-circUNC13A nor the UNC13A cryptic exon is detected. **I)** Correlation between ASO-1-mediated reductions in c-circUNC13A expression and restoration of correctly spliced UNC13A transcript. Data points are coloured according to the ASO-1 concentration Grey bars indicate the mean ± SD expression levels in cells without ASO treatment. **J)** Schematic representation of the in vivo experimental design to evaluate the effect of UNC13A cryptic exon-targeting ASO treatment on c-circUNC13A levels. BAC transgenic mice expressing the entire human UNC13A received intracerebroventricular (ICV) administration of the ASO, followed by intravenous administration of an AAV expressing a miRNA to induce TDP-43 knockdown **K)** RT– qPCR analysis of c-circUNC13A expression in the cortex, brainstem, caudal spinal cord (SC) and rostral spinal cord (SC) of human UNC13A BAC transgenic mice following AAV-mediated TDP-43 knockdown, with or without treatment with the UNC13A cryptic exon-targeting ASO. c-circUNC13A expression was normalised to TBP mRNA and is presented as percent expression relative to the TDP-43 knockdown condition, set to 100%. Each dot represents an individual mouse. Statistical significance was assessed using a one-way Anova with correction for multiple test comparison (n.s. p > 0.05, ***p < 0.001).

We then asked whether the UNC13A CE and c-circRNA have different stability. We took advantage of an engineered i3Ns line where endogenous TDP-43 can be rapidly degraded through addition of PROTAC^14^, and investigated UNC13A CE and c-circRNA at different time-points after TDP-43 depletion. UNC13A CE levels rapidly rise and plateau 24 hours after treatment. In contrast, *c-circUNC13A* displayed a slower and more progressive increase throughout the time course, continuing to accumulate over 72 hours, (**Fig. 4C, left panel**). We then compared how UNC13A CE and c-circRNA persist when TDP-43 levels are restored after 14 days of TDP-43 depletion PROTAC - this was achieved by replacing PROTAC with a catalytically inactive competitor (PROTACi). UNC13A CE levels declined rapidly, reaching approximately 40% of their initial abundance within 24 hours, whilst *c-circUNC13A* showed markedly slower decay kinetics, compatible with a longer half-life, retaining 80% of its initial levels at the same time point and decreasing gradually thereafter (**Fig. 4C, right panel**). These findings indicate that *c-circUNC13A* is substantially more stable than the corresponding linear cryptic transcript.

### c-circUNC13A and UNC13A CE are co-regulated

Clinical trials aimed at correcting UNC13A cryptic exon inclusion are under way^13,14^. As c-circUNC13A and UNC13A CE are both non-functional alternatives to the correct exon 20-21 splicing (**Fig. S9B**), it is paramount to understand whether they are co-regulated, or whether suppressing UNC13A CE results in c-circUNC13A increase.

We therefore analysed SHSY-5Y cells where the UNC13A CE region was removed by CRISPR/Cas9-mediated genome editing (ΔCE) (**Fig. 4D**)^14^. As previously reported, UNC13A CE was not produced in the ΔCE line upon TDP-43 depletion. We also found c-circUNC13A induction upon TDP-43 knock-down in the ΔCE line to also be completely abolished, demonstrating that the region producing the cryptic exon is essential for the expression of the circular isoform (**Fig. 4E**).

An important point to address is whether the suppression of c-circUNC13A in the ΔCE line is due to the deletion of part of intron 20 sequence or it’s dependent on the UNC13A CE formation being suppressed. We therefore investigated the effect of different splice-switching ASOs that reduce the UNC13A CE on c-circUNC13A^14^. Whilst non-targeting ASOs displayed no effect on UNC13A CE and c-circUNC13A, UNC13A CE ASOs led to the reduction of up to 80% of the circRNA levels (**Fig. S12A**). We then used different doses of a high-performing ASO (**Fig. 4F**) and observed a dose-dependent reduction effect on the circRNA (**Fig. 4G**), which also displayed positive correlation with UNC13A CE (**Fig. 4H**). Importantly, c-circUNC13A also showed a negative correlation with the correctly spliced UNC13A isoform (**Fig. 4I**), supporting c-circUNC13A levels as a potential readout of UNC13A rescue.

We next tested whether the coordinated regulation of UNC13A cryptic exon inclusion and c-circUNC13A formation also occurs in vivo. Since UNC13A CE is not conserved in rodents, we used a BAC transgenic mouse expressing the entire human UNC13A locus (**Fig. 4J**). Following TDP-43 knockdown by an AAV-delivered Tardbp miRNA, c-circUNC13A became readily detectable in the CNS by RT-qPCR. Intracerebroventricular injection of an ASO targeting the UNC13A CE suppressed c-circUNC13A formation (**Fig. 4K**), demonstrating that therapeutic modulation of UNC13A mis-splicing is coupled by reduction of the c-circRNA in vivo.

Overall, these results show that circUNC13A is more stable and detectable than UNC13A CE, that the biogenesis of the two events is linked, and highlights the potential of c-circUNC13A as a target engagement biomarker for therapies modulating UNC13A CE.

## Discussion

We identify c-circRNAs as a novel consequence of TDP-43 dysfunction. We show that loss of nuclear TDP-43 induces widespread alterations in back-splicing and formation of de novo circRNAs that are absent under physiological conditions. These c-circRNAs are specifically associated with ALS and FTD cases with TDP-43 pathology, display increased stability and superior detectability compared with their associated linear cryptic counterparts, and can reveal previously unappreciated layers of RNA misprocessing. Finally, we demonstrate that c-circUNC13A is mechanistically linked to the disease-relevant UNC13A CE and may serve as a target-engagement biomarker for emerging therapies directed at correcting *UNC13A* splicing.

While TDP-43 has long been recognised as a repressor of canonical splicing^7,10,11^, our findings identify an additional role for this RNA-binding protein in circRNA biology. Beyond regulating the abundance of physiological circRNAs, TDP-43 acts to suppress the formation of cryptic circRNAs (c-circRNAs). The enrichment of TDP-43 binding sites within the introns flanking c-circRNAs supports a direct role for TDP-43 in preventing aberrant back-splicing at these loci. Although the factors driving cryptic back-splicing remain to be elucidated, widespread defects in RNA processing observed once TDP-43 is removed, including cryptic splicing and intron retention^24^, suggest that its depletion creates a permissive environment for aberrant back-splicing. Perturbations in splicing kinetics may favour back-splicing, which is generally considered less efficient than linear splicing^38^, by increasing the time window during which non-canonical splicing sites can pair^39^. Since the back-splicing reaction relies on the formation of a loop which brings together the two introns flanking the circRNA-producing exons^27,28^, one possibility is that loss of TDP-43 binding exposes intronic low-complexity sequences that become available for intermolecular RNA interactions^40^, thereby promoting circularisation.

The discovery of TDP-43-dependent cryptic events (CEs)^4–7^ has improved our understanding of ALS and FTD pathogenesis by providing specific molecular readouts of TDP-43 loss of function^10,11^. In practice, however, several obstacles limit their utility as biomarkers. Many CE-containing transcripts are rapidly degraded through nonsense-mediated decay^15,35,41^, while those that escape degradation remain intrinsically unstable and are particularly vulnerable to degradation in biofluids^42^. We and others have discovered that a fraction of CEs encodes for additional in-frame peptide sequences and therefore give rise to novel proteins^8,9^. Although current assays for cryptic peptides are promising, they still require sensitivity improvements for clinical applications. On the other hand, circRNAs possess properties that make them particularly attractive biomarker candidates. Their covalently closed structure provides them with exceptional stability^43,44^, and, despite representing only a minor fraction of cellular RNAs, circRNAs are highly enriched in biofluids and extracellular vesicles^45^ due to their resistance to exonucleases. In addition, their circular structure enables selective signal amplification through rolling-circle reverse transcription^31^, providing a technical advantage over linear cryptic transcripts, which are frequently undetectable in tissue because of low abundance and rapid degradation^15,35,41^.

The applicability of c-circRNAs as potential biomarkers is exemplified by the lead candidates that we have identified (*c-circUNC13A*, *c-circBCAR1*, *c-circDIP2C* and *c-circDSCAM*). These were highly enriched in both ALS-TDP and FTD-TDP post-mortem CNS samples and showed excellent discrimination between TDP-43 and non-TDP proteinopathies. Using a targeted rolling-circle amplification approach, these molecules achieved near-complete separation of disease groups, with the combined model reaching an AUC of 0.99. Together, these findings establish c-circRNAs as highly sensitive and specific molecular footprints of TDP-43 dysfunction and support their development as biomarkers for TDP-43 proteinopathies.

An unexpected observation was the frequent coexistence of c-circRNAs with other forms of aberrant splicing within the same genes. Approximately one-fifth of High-Confidence c-circRNAs were associated with neighbouring cryptic or skiptic exons, suggesting that certain transcripts are particularly vulnerable to multiple forms of RNA misprocessing following TDP-43 depletion. circRNA biogenesis has already been reported to co-occur with non-canonical splicing events. For example, depletion of the splicing factor SFPQ leads to widespread alterations in circRNA biogenesis accompanied by upstream intron retention and cryptic splicing events^39^. More recently, depletion of HNRNPM has been shown to promote both aberrant exon inclusion and circRNA biogenesis, further supporting the idea that impaired splice site selection creates a permissive environment for back-splicing^46^.

Among the genes displaying cryptic multi-misprocessing events we identified *UNC13A*, *RPTOR* and *EHMT1*. All these transcripts undergo aberrant splicing events that generate premature termination codons and trigger NMD-mediated transcript degradation. While the consequences of *UNC13A* missplicing have been extensively characterised^14,24^, our work provides the first evidence that TDP-43 dysfunction also leads to loss of RPTOR and EHMT1 proteins through cryptic RNA processing mechanisms. Importantly, dysfunction of both proteins has previously been implicated in neurological disease. RPTOR is an essential component of the mTORC1 complex and its reduction impairs mTOR signalling, promotes TFEB activation and alters autophagy-lysosomal pathways that have been implicated in neurodegeneration^22,23^. Similarly, EHMT1 is a key histone methyltransferase required for neuronal development and synaptic function, and haploinsufficiency or mutation of EHMT1 causes Kleefstra syndrome, a severe neurodevelopmental disorder characterised by intellectual disability and cognitive impairment^36,37^. The reduction of these proteins suggests TDP-43 could contribute to neurodegeneration through additional pathogenic pathways, some of which remain hidden because the underlying cryptic transcripts are rapidly degraded.

Importantly, the same cryptic RNA processing events that reduce the abundance of disease-relevant proteins also generate stable c-circRNAs that remain readily detectable despite degradation of the corresponding linear transcripts. This difference is also reflected in their subcellular localisation. While cryptic linear transcripts (CEs) subject to NMD are predominantly found within the nucleus^15^, c-circRNAs accumulate in the cytoplasm, where they are more likely to persist and potentially be incorporated into extracellular vesicles, structures where circRNAs have already been shown to be enriched^45^. Coupled with their resistance to NMD and selective amplification by rolling-circle reverse transcription, these features contribute to their superior detectability in patient-derived samples. Therefore, c-circRNAs can serve as stable molecular footprints of cryptic splicing events that would otherwise remain largely invisible in human tissue. Although the evaluation of c-circRNAs in extracellular vesicles and biofluids was beyond the scope of the present study, our findings provide a strong rationale and a set of candidate molecules for future studies aimed at assessing their potential as EV-associated biomarkers of TDP-43 dysfunction.

Among the identified c-circRNAs, *c-circUNC13A* is a peculiar case because it arises from one of the strongest genetic risk loci for both ALS and FTD^24,25,32–34^ and its linear counterpart is currently being targeted for therapeutic intervention^13,14^. Multiple orthogonal lines of evidence demonstrate that *c-circUNC13A* and the disease-linked *UNC13A* cryptic exon are mechanistically linked. Deletion of the genomic region responsible for cryptic exon formation abolished induction of both transcripts, while splice-switching antisense oligonucleotides suppressed both species in a dose-dependent manner. These observations indicate that *c-circUNC13A* is not an independent by-product of TDP-43 dysfunction but rather a coordinated consequence of the same pathogenic missplicing event.

*c-circUNC13A* possesses several properties that make it an attractive target-engagement biomarker. It is considerably more stable than the corresponding linear cryptic transcript as it accumulates progressively following TDP-43 depletion and persists longer after TDP-43 function is recovered. These kinetic differences suggest that *c-circUNC13A* may act as a time-averaged molecular readout of TDP-43 dysfunction, providing a more durable molecular footprint of pathology. This could buffer transient fluctuations in splicing activity and therefore be particularly valuable in therapeutic settings to monitor treatment response.

Collectively, our findings extend the paradigm of TDP-43-dependent RNA pathology beyond linear missplicing and identify cryptic circular RNAs as previously unrecognized consequences of nuclear TDP-43 loss. By combining disease specificity, molecular stability and enhanced detectability, these molecules represent a promising new class of biomarkers and provide novel insights into the complex RNA processing defects that underlie TDP-43-dependent ALS and FTD pathology. Furthermore, our work establishes a framework to support the future evaluation of cryptic circular RNAs in extracellular vesicles and other biofluids, both as biomarkers of TDP-43 pathology and as pharmacodynamic readouts of therapeutic intervention.

## Materials and Methods

### Human induced pluripotent stem cell (iPSC) culture

In this study we used the WCT11 iPSC that were previously engineered to overexpress human neurogenin-2 (NGN2) to generate cortical-like i3Neurons, or an hNIL construct overexpressing human transcription factors Islet-1 (ISL1), LIM Homeobox 3 (LHX3), and NGN2, to generate i3-lower motor neurons (i3-LMNs), both under a doxycycline-inducible promoter, alongside an expression cassette for an enzymatically dead Cas9 (+/− CAG-dCas9BFP-KRAB), as previously described. These constructs were integrated into the AAVS1 and CLYBL safe harbour loci, respectively47,48. The generation of the Halo-TDP line is described elsewhere14. iPSCs were maintained in Essential 8 Flex medium (Gibco, A2858501) with daily feeds and kept in an incubator at 37°C and 5% CO2. Cells were routinely passed using Versene (Gibco, 15040066) in Geltrex-coated (1:100; Gibco, A14133-01) tissue culture dishes. STR profiling and mycoplasma contamination test were performed periodically.

### Differentiation of iPSCs into cortical-like neurons (i3Neurons) and culture

The differentiation process was as reported previously^49^. To initiate neuronal differentiation, iPSCs were dissociated using Accutase (Gibco, A1110501) and re-plated onto Geltrex-coated (1:100; Gibco, A14133-01) tissue culture dishes in induction medium: DMEM/F-12 + GlutaMAX medium (Gibco, 31331028), 1x MEM Non-Essential Amino Acids (Gibco, 1140050), 2 μg/ml Doxycycline hyclate (SigmaAldrich, D9891), 1x N2 supplement (Gibco, 17502048), 2 μM XAV939 (Cambridge Bioscience, SM38-10), 10 μM SB431542 (Biotechne, 1614), and 100 nM LDN-193189 (Cambridge Bioscience, 19396-5mg-CAY). 1x ROCK Inhibitor Y-27632 dihydrochloride (Tocris, 1254/10) was added to the medium on the day of plating. Induction medium was changed daily during this stage. On day 3, pre-neuron cells were re-plated onto dishes coated with 50 μg/ml Poly-D-Lysine (Gibco, A38904-01) and 10 μg/ml Laminin (Gibco, 23017015) in 12-well dishes (500,000 cells per well) for RNA and protein extraction. They were plated in i3Neuron cortical neuron culture medium: BrainPhys neuronal culture medium (StemCell Technologies, 5790), supplemented with 1x N21 Max (R&D System,AR008), 1x N2 Max (R&D System,AR009), 10 ng/ml BDNF (PeproTech, 450-02), 10 ng/ml GDNF (PeproTech, 450-10), and 1 μg/ml Laminin (Gibco, 23017015). To facilitate single-cell survival, 1x ROCK Inhibitor was added on the day of plating. 24 hours after plating, culture medium was fully replaced to remove the ROCK Inhibitor and add CultureOne supplement (antimitotic agent; 1:100;Gibco, A3320201). Cells were kept in i3Neuron cortical neuron culture medium for a variable period of time, 14-30 days for the CRISPRi-mediated TDP-43 depletion and 28 days for the PROTAC-mediated TDP-43 depletion (see below).

### Differentiation of iPSC-derived lower motor neurons (i3LMNs)

Differentiation of human iPSCs was performed as previously reported. Briefly, iPSC induction was achieved using induction medium containing DMEM/F12, GlutaMAX supplement medium (Thermo Fisher), MEM non-essential amino acids (Thermo Fisher), N2 supplement (Thermo Fisher), 0.2 mM compound E, 2 μg/ml doxycycline, Pen/Strep and 1 μg/ml Laminin. After induction for 48 h, cells were dissociated from plates by Accutase and reseeded on poly-D-lysine (PDL)/laminin-coated tissue culture dishes in induction medium supplemented with 1x ROCK Inhibitor and CultureOne supplement. After 24 h, induction medium was replaced by motor neuron medium containing Neurobasal (Gibco, 21103049), Glutamax supplement medium (Thermo Fisher), MEM non-essential amino acids (Thermo Fisher), N2 max supplement (R&D Systems, AR009), N21 max supplement (R&D Systems, AR008), CultureOne supplement, 1 μg/ml Laminin, 2 μg/ml Doxycycline hyclate, 10 ng/mL BDNF (Peprotech) and 10 ng/ml GDNF (Peprotech). Half-medium changes were performed twice a week.

### TDP-43 knockdown in SH-SY5Y and SK-N-BE(2) cells

SH-SY5Y and SK-N-BE(2) cells were transduced with SmartVector lentivirus (V3IHSHEG_6494503) containing a doxycycline-inducible shRNA cassette for TDP-43. Transduced cells were selected with puromycin (1 μg ml−1) for one week. The pool of TDP-43-knockdown SH-SY5Y and SK-N-BE(2) cells was plated as single cells and expanded to obtain a clonal population. The generation of SH-SY5Y cell line with UNC13A cryptic exon deletion (ΔCE line) was described in Keuss 202414. Cells were grown in DMEM/F12 containing Glutamax (Gibco) supplemented with 10% FBS (Gibco) and 1% Penicillin/Streptomycin (Thermo Fisher). For induction of shRNA against TARDBP, cells were treated for 10 days with increasing amounts of doxycycline hyclate (1000 ng/ml) (Sigma), unless differently specified.

### CRISPRi-mediated TDP-43 knockdown in human iPSCs

To achieve knockdown, sgRNAs targeting either TARDBP/TDP-43 or a non-targeting control guide were delivered to iPSC by lentiviral transduction. To produce the virus, human embryonic kidney (HEK) cells were transfected with the sgRNA, packaging (pCMVΔ8.91) and VSVG envelope (pMD.G) plasmids usingTrans-IT 293 (Cambridge Bioscience, cat. No. MIR2700), then cultured for 2–3 days in the following media: opti-MEM Reduced Serum + Glutamax (Thermo Fisher Scientific, cat. no. 51985-026) with 5% FBS (Sigma, cat. no. TMS-013-B). After media filtration, virus was concentrated from the media 1:10 in PBS using Lenti-X concentrator (Takara Bio, cat. no. 631231), aliquoted and stored at −80 °C for future use. The sgRNAs were cloned into pU6-sgRNA EF1Alpha-puro-T2A-BFP vector12,37 (gift from J. Weissman; Addgene 60955)47,50. sgRNA sequences were as follows: non-targeting control: GTCCACCCTTATCTAGGCTA, TARDBP: GGGAAGTCAGCCGTGAGACC (strong guide). Virus was delivered to iPS cells in suspension following an Accutase split. After 24 hours, cells were washed with PBS and fed with fresh E8 media. Two days after lentiviral delivery, cells were selected for 48h with Puromycin (10 μg/ml). iPSCs were then expanded 1–2 days before initiating neuronal differentiation. Knockdown efficiency is tested by qPCR and western blotting.

### PROTAC-mediated TDP-43 degradation and recovery

For this experiment, 500k i3 HALO-TDP neurons were plated in 12 well plates previously coated with with 50 μg/ml Poly-D-Lysine (Gibco, A38904-01) and 10 μg/ml Laminin (Gibco, 23017015). The neurons were maintained in i3Neuron cortical neuron culture medium: BrainPhys neuronal culture medium (StemCell Technologies, 5790), supplemented with 1x N21 Max (R&D System,AR008), 1x N2 Max (R&D System,AR009), 10 ng/ml BDNF (PeproTech, 450-02), 10 ng/ml GDNF (PeproTech, 450-10), and 1 μg/ml Laminin (Gibco, 23017015), with half media change performed twice a week. To induce TDP-43 knockdown, Halo PROTAC-E 300 nM (Division of Signal Transduction Therapy DSTT^51^, was added to cultures with each media change starting on DIV14. To restore TDP43 protein expression following knockdown, the neurons were treated with Halo PROTAC-E inactive epimer 300 nM (DSTT) at defined timepoints prior to harvesting at DIV28.

### iPSC maintenance and differentiation (for ASO treatment)

The GM25256 human iPSC line (Coriell Institute) was maintained in mTeSR1media (StemCell Technologies) on Matrigel (Stemcell Technologies, Corning). Human iPSCs were fed daily and split every 4-5 days using ReLeSR (StemCell Technologies) according to the manufacturer’s instructions. Differentiation of iPSCs into motor neurons was carried out using 3D developmental pathways for 14 days (Maury, 2015; Ma, 2022). On Day 14, neuronal spheroids were dissociated and plated on Poly-D-Lysine/Laminin coated 96-well plates (Phenoplate, Revvity) and maintained in Neurobasal Medium (Thermo Fisher) containing neurotrophic factors (BDNF, GDNF and CNTF; R&D Systems). Cells were treated with 5-Fluorouracil 4 days post plating to remove cycling progenitors. Rat astrocytes (Stem Cell Technologies) were added once to the motor neuron cultures 8-17 days after plating.

### ASO Preparation

To make 100 µg/µL storage stock solution, lyophilized ASO (TDP-43 gapmer or UNC13A splice correctors) was resuspended in the appropriate amount of sterile PBS, pH 7.4. Once fully dissolved, the ASO was filter sterilized using a spin column (0.22 µM). The 100 µg/µL storage stock solution of ASO was aliquoted and stored at –20°C for future use.

To make ∼500 µM working stock solutions for dosing neurons, the 100 µg/µL storage stock solution was diluted appropriately with sterile PBS, pH 7.4 and filter sterilized using a spin column (0.22 µM). To determine the concentration of the ASO working stock solution, the absorbance at 260 nm was measured on a NanoDrop One (ThermoFisher Scientific) and concentrations were calculated using the molecular extinction coefficient for that ASO. The working stock solution of ASO was aliquoted and stored at –20°C for future use. Dilutions of working stocks to the appropriate dose levels were made with the appropriate amount of sterile PBS, pH 7.4.

### TDP-43 KD and antisense oligonucleotide (ASO) treatment in cells

On Day 19-21 after neuronal plating, TDP-43 gapmer (2 μM) or equivalent volume of PBS (NT) was added once to cells through gymnotic (free) uptake to knockdown (KD) TDP-43. On Day 22-27 after neuronal plating, splice correcting ASOs or control ASO (nusinersen sequence) were added once to cells through gymnotic uptake. TDP-43 gapmer (at 2 µM) or equivalent PBS control was subsequently added to cells through gymnotic uptake once a week for the following 3 weeks to maintain consistent KD levels. After at least 40 days in culture from the time they were plated, motor neurons were processed to assess RNA transcript levels by RT-qPCR.

RT-qPCR was performed using the TaqMan Fast Advanced Cells-to-CT kit (Thermo Fisher Scientific) according to the manufacturer’s instruction.

All assays were purchased from Thermo Fisher Scientific. Percent expression of correctly spliced UNC13A mRNA was calculated using the ΔΔCT method and normalized to control (NT) treated cells. Percent expression of UNC13A cryptic exon mRNA and c-circUNC13A was calculated using the ΔΔCT method and normalized to TDP-43 KD only cells. Undefined CT values for UNC13A cryptic exon and c-circUNC13A (such as in control samples) were converted to 0% expression. All analyses were performed in R. All sequences are made of fully 2’MOE modified bases with PS linkages unless otherwise specified, i.e the gapmer has DNA bases as well.

### Human UNC13A BAC transgenic mice

hUNC13A BAC Tg mice were generated at Taconic Biosciences (Germantown, NY) by pronuclear injection of a BAC subclone carrying the human UNC13A (hUNC13A) gene into a mouse embryo. Potential founder lines were screened for neuronal specific hUNC13A expression, with a single founder used to establish the colony. Transgene mapping determined that these mice possess two copies of the hUNC13A gene in tandem and the integration did not occur within any known protein coding gene. Note, both copies of the endogenous mouse Unc13a gene (Munc13-1) are still present in these mice. Mice were maintained as a heterozygous colony for the hUNC13A allele. No abnormal phenotypes have been documented for this mouse line. Female hUNC13A mice aged 8-12 weeks at experiment start were used for these studies.

### Intracerebroventricular administration of antisense oligonucleotides

For ICV injections, adult mice were administered peri-operative analgesic and placed in a stereotaxic frame under continuous isoflurane anesthesia. Their ears were secured using ear bars and a nose cone after which fur was removed and the skin disinfected. A midline incision was then made along the top of the skull, and the skin was retracted to expose the bregma and lambda bone suture junctions. The skull was leveled by aligning bregma and lambda, and the injection site was identified (−0.5 anterior-posterior, +1.0 medial-lateral). Using the stereotaxic arm a small burr hole was drilled above the injection site using a sterile bit. A 100 μl Hamilton microsyringe was lowered through the craniotomy site so that the bevel opening was −1.75 mm dorsal-ventral below the brain surface. Ten microliters of 30 mg/mL ASO 6 or aCSF solution was injected using an automated UMP3 UltraMicroPump. The needle was left in place for 3 minutes and then withdrawn, and the scalp incision was sutured. The mice were allowed to recover from anesthesia in a heated recovery cage and then returned to their home cage.

### rAAV-mediated Tardbp knockdown

hUNC13A BAC Tg mice were injected 1 week post ICV surgery in the lateral tail vein with 100-200 μL total volume rAAV containing a miRNA sequence targeting Tardbp. The rAAV was packaged by Packgene Biotech, Inc. (Houston, TX), with vector titers determined at Trace Neuroscience using payload specific primer/probe sets. Animals were injected with rAAV based on vector genomes per kilogram mouse body weight. This rAAV utilizes the PHP.eB capsid and C57BL/6 background of the hUNC13A BAC mice to enable widespread neuronal transduction with intravenous injection.

### Mouse tissue harvesting and RT-qPCR

Mice were euthanized 4 weeks after ICV surgery (3 weeks post rAAV administration) by anesthesia with isoflurane and decapitation. Brain and spinal cord tissues were harvested, placed into soft tissue homogenizing tubes (Bertin, P000912-LYSK0-A or P000933-LYSK0-A), weighed, and flash frozen in liquid nitrogen. Tissues were homogenized on a Precellys Evolution Touch homogenizer (Bertin Technologies) in lysis buffer from the MagMax mirVana Total RNA Isolation Kit (Thermo Fisher Scientific). RNA was extracted using the bead-based MagMax mirVana Total RNA Isolation Kit (Thermo Fisher Scientific) with a KingFisher Apex instrument (Thermo Fisher Scientific). RNA was quantified via Qubit Fluorometer (Thermo Fisher Scientific) and the same amount of RNA per tissue for each mouse was reverse transcribed to cDNA using SuperScript IV VILO Master Mix (Thermo Fisher Scientific). Taqman qPCR was performed using TaqMan Fast Advanced Master Mix (Thermo Fisher Scientific) according to the manufacturer’s instructions using the following multiplexed assays: c-circUNC13A and Hprt1 as the housekeeper. Both assays were purchased from Thermo Fisher Scientific. Percent expression of c-circUNC13A was calculated using the ΔΔCT method with normalization to Hprt1 and normalization to TDP-43 KD/aCSF ICV treatment conditions.

### RNA isolation and analysis

RNA isolation was carried out using either the RNeasy® mini kit (Qiagen) or the Direct-zol RNA Miniprep (Zymo research), according to manufacturer’s protocol with an optional on-column DNase I digestion step. RNA concentrations were measured by NanoDrop One (ThermoFisher Scientific), and equal quantities – unless differently specified - were used for reverse transcription.

Reverse transcription was performed using either RevertAid cDNA synthesis kit (ThermoFisher Scientific), Superscript IV (ThermoFisher Scientific) or Induro (NEB). For RevertAid and Superscript IV, the reaction was carried out according to the manufacturer’s protocol. For Induro, RNA denaturation and primer annealing was performed by incubating the template RNA with 1 μL of 100 μM random hexamers in a volume of 12 μL at 65°C for 5 minutes, followed by cooling on ice for 2 minutes. A mixture containing 4 μL of 5 X Induro RT Reaction Buffer (NEB), 2 μL MgCL2 30mM (Sigma-Aldrich), 1 μL 10 mM dNTPs (ThermoFisher Scientific), 1 μL RiboLock RNase Inhibitor 20U/ μL (ThermoFisher Scientific) and 200 U of Induro RT (NEB) was then added and the reaction was incubated at 23°C for 10 minutes, at 30°C for 5 minutes and at 55°C for 1 hour, followed heat-inactivation at 95°C for 1 minute.

RT-qPCR analyses were performed using PowerUp SYBR Green Master Mix reagent (ThermoFisher Scientific) or TaqMan™ Multiplex Master Mix (ThermoFisher Scientific). DNA amplification was monitored on a QuantStudio 5 Real-Time PCR system (Applied Biosystems) using the ΔΔCt method. Relative RNA quantity was calculated as the fold change (2−ΔΔCt) with respect to the experimental control sample set as 1 and normalized to a housekeeping gene (GAPDH mRNA or TDP mRNA), unless differently specified.

### Receiver operating characteristic analyses

Receiver operating characteristic (ROC) curve analyses were performed on quantitative PCR data consisting of Ct values normalized to GAPDH reference control (ΔCt), which were subsequently log10-transformed prior to analysis. For each individual circRNA, ROC curves were generated by comparing the log10-transformed normalized expression values between FTD-TDP cases and the combined control group (CTRL and FTD-non-TDP). The area under the ROC curve (AUC) and its 95% confidence interval (CI) were calculated using the R package pROC with default parameters, where CIs were estimated using DeLong’s method. A combined classifier was generated by fitting a multivariable logistic regression model including all circRNA markers as predictors and FTD-TDP status as the binary outcome. Predicted probabilities from the logistic regression model were subsequently used to generate a ROC curve and calculate the corresponding AUC for the combined classifier.

### Comparison between Induro and Superscript IV activity

For the comparison between Superscript IV and Induro, 500 ng of an in vitro transcribed RNA carrying c-circUNC13A nucleotide sequence – either in a circular or linearised version preserving the back-splicing junction sequence – were reverse-transcribed with either Superscript IV (200 U) or Induro (200 U) in a 20 μL reaction with random hexamers, following manufacturer’s instructions for Superscript IV and the aforementioned protocol for Induro. After completion of the reactions, 1 μL from each sample was diluted and used for RT-qPCR analysis. The rest was first treated with Proteinase K (ThermoFisher Scientific) and incubated at 37°C for 15 minutes and at 55°C for 20 minutes. The reactions were then purified with Mag-Bind® TotalPure NGS Beads (Omega Bio-tek) according to the manufacturer’s protocol and eluted in 26 μL water. A fraction of each sample was saved, while the rest was treated with 1,5 μL RNase H (NEB) and 1.5 μL RNase If (NEB), incubated at 37°C for 30 minutes and at 70°C for 20 minutes. All samples – either before or after RNase treatment – were run on a 0.8% TAE agarose gel next to 1 kb Plus DNA Ladder (NEB) and then stained with SYBR™ Gold Nucleic Acid Gel Stain (ThermoFisher Scientific) before visualization.

### RNase R treatment

For RNase R treatment, 3 μg of total RNA [DD1] were diluted in 20 μL reaction with 5 U of RNase R (Lucigen), then incubated 15 min at 37 °C. The reactions were purified by adding 300 μLTRIzol (ThermoFisher Scientific) supplemented with 4 pg of a DNA spike-in molecule, later used for RT-qPCR normalization, and by proceeding with phenol-chloroform extraction.

### Nonsense-mediated decay (NMD) inhibition and Subcellular fractionation

For the fractionation experiments, SH-SY5Y cells were treated for 10 days with 25 ng/ml doxycycline hyclate (Sigma) to induce the shRNA against TDP-43, and for 24 hours with 0.5 μM SMG1-11j inhibitor^52^ to block the NMD machinery. After 10 days, cells were trypsinised, pelleted and resuspended 467 in 1X PBS (Thermo). A fraction of the resuspended cells was pelleted and used for the subcellular fractionation with the Ambion PARIS Kit (Life Technologies), according to the manufacturer’s instructions. RNA from the nuclear and the cytosolic fractions was extracted with the Direct-zol kit (Zymo) with on-column DNase I treatment and quantified by NanoDrop One (ThermoFisher Scientific). For each experimental condition, 2 μg of cytoplasmic RNA and an equal volume of nuclear RNA fraction were reverse-transcribed with RevertAid First Strand cDNA Synthesis Kit (Thermo Fisher Scientific) according to the manufacturer’s instructions. RT-qPCR was performed with PowerUp SYBR Green Master Mix reagent (ThermoFisher Scientific). DNA amplification was monitored on a QuantStudio 5 Real-Time PCR system (Applied Biosystems) The relative abundance of each transcript in the nuclear and cytoplasmic fractions was estimated from Ct values as 2^−Ct, and the proportion of transcript in each compartment was calculated relative to the sum of the two fractions, which was defined as 100%.

### Western blotting

i3Neurons were homogenized in RIPA lysis buffer (Thermo Fisher Scientific, cat. no. 89900) supplemented with cOmpleteTM EDTA-free protease inhibitor cocktail (Roche, cat. no. 11836170001) and PhoSTOP Phosphatase inhibitor (Roche, 4906845001). Samples were loaded on a NuPAGE 4−12% Bis-Tris protein gel (Invitrogen) and run in NuPAGE MOPS buffer. Proteins were transferred onto the nitrocellulose blotting membrane using the Trans-Blot Turbo 5x Transfer Buffer (BioRad, 10026938) using the Trans-Blot Turbo Transfer System for 30 minutes. The membrane was blocked in 5% non-fat milk in TBST (20 mM Tris, 150 mM NaCl and 0.1% Tween 20) and probed overnight with primary antibodies diluted in 5% non-fat milk in TBST (anti-RPTOR (Proteintech, 20984-1-AP) 1:2000, anti-EHMT1 (Bethyl Laboratories A301-642A) 1:2000, anti-TDP-43 (Proteintech,10782-2-AP) 1:2,000 and anti-TUBULIN (Proteintech,66031) 1:2,000). After 1-hour incubation with horseradish peroxidase (HRP)-conjugated secondary antibodies diluted in 5% milk in TBST (anti-mouse HRP (Bio-Rad, 1706516) 1:10,000 and anti-rabbit HRP (Bio-Rad, 1706515) 1:10,000), the membrane was developed using Immobilon Classico HRP substrate (Sigma-Aldrich) and the Bio-Rad ChemiDoc system.

### RNA-sequencing and pre-processing

SH-SY5Y, SK-N-BE(2), and i3N sequencing libraries were prepared with KAPA RNA HyperPrep with RiboErase Kit and sequenced on a NovaSeq 6000 machine. SH-SY5Y were paired-end 150 bp and i3N were paired-end 100 bp.

Samples are quality trimmed using Fastp with the parameter “qualified_quality_phred: 10”, and aligned to the GRCh38 genome built using STAR^53^ (v2.7.0f) with gene models from GENCODE v31. Gene expression is quantified using FeatureCounts using gene models from GENCODE v31.

### CircRNA detection with CIRI2

Processed reads are aligned to the human reference genome (GRCh38) using BWA-MEM^54^ (v0.7.17) with -T 19 option. CircRNA detection in each sample is then carried out using CIRI2^26^ software (v2.0.6), which is able to identify circRNAs by searching for reads that map to back-splicing junctions. To identify circRNA host genes, the program is provided with Ensembl 108 gene annotation.

For each back-splicing event found, CIRI2 reports the number of reads mapping to the back-splicing junction and on the corresponding linear splicing junctions, calculated summing all the reads mapping linearly on both the splice junctions involved in back-splicing; the latter are not reported if no read is assigned to the back-splicing junction, even if the circular RNA is detected in other samples. In order to count the reads mapping to linear splicing sites (cognate linear splicing junction, SJ) in samples in which no reads were mapped to corresponding back-splicing junctions detected in other samples, alignment files from each sample were modified by adding reads mapping to circRNAs found only in other samples and CIRI2 was rerun on these files. Alignments files operations were performed using Picard suite^55^ (v2.24.1) (https://broadinstitute.github.io/picard/) and SAMtools^56^ (v1.10) (http://www.htslib.org).

### Differential Expression Analysis

To evaluate the differential expression of circRNAs between knock-down and control conditions, the edgeR^57^ R package (v3.34.1) is provided with the read counts of the back-splicing events detected. Low counts events are not tested for differential expression. The low counts threshold has been settled based on the number of samples per condition, specifically: 2 counts in at least 2 samples for SH-SY5Y and SK-N-BE(2), 2 counts in at least 5 samples for i3LMN and 2 counts in at least 6 samples for i3N.

For each contrast, samples are normalized using standard edgeR normalization (TMM); model fitting and testing is performed using the glmFIT and glmLRT functions.

Reads mapping to back-splicing junctions are converted to Count Per Million (CPM) values using edgeR and used for circRNA quantification.

Given the low number of reads used for testing, we decided to use p-value instead of false discovery rate to select for differentially expressed events, setting the significance threshold value to 0.05.

We identified as “High-Confidence” the circRNAs significantly upregulated in at least 2 out of 4 cell lines. We considered “cryptic” the upregulated in TDP-KD condition, with less than 1 count per control sample.

For the analyses of linear RNAs and of the cognate linear splicing junction we applied the same counts thresholds.

Heatmaps and Upset plot graphical representations were depicted using ComplexHeatmap^58^ R package (v2.8.0).

### CLR analysis

For each sample, the circular back splicing junction counts (BSJ) and the cognate linear splicing junction counts (SJ) were extracted from the CIRI2 output file. For each circRNA event, circular-to-linear ratio (CLR) was computed. CLR was calculated as:

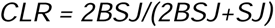

Following the CIRI2 manual, the BSJ counts were multiplied by two to correct the asymmetry between circular and linear read contributions: each BSJ-spanning read represents sequence originating from both sides of the circular junction but is counted only once, whereas linear (non-junction) reads derive from a single end.

A mean CLR was then calculated separately for the CTRL and TDP-43 knockdown (KD) groups, and the difference between group means (ΔCLR) was used to quantify changes in circularization:

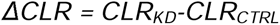

To assess statistical significance, we applied a binomial generalized linear model (GLM) in which 2BSJ values were treated as “successes” and SJ values as “failures.” For each circRNA event, two nested models were fitted: a null model assuming no effect of condition (CLR constant across samples) and a full model including condition as a predictor. A likelihood ratio test (LRT) was then used to compare the models and determine whether incorporating the condition term significantly improved model fit.

### TDP-binding sites analysis

TDP binding regions were retrieved from POSTAR3 database^29^. Upregulated circRNAs (candidates) were tested for TDP binding in their flanking introns (specifically, in the +-1000bp region from the BSJ, or less if intersecting with an exonic region).

For each candidate, 2 control circRNAs were selected from invariant circRNAs (p-value>0.05) taking into consideration the properties of the flanking introns (length, %GC, distance between back-splicing exons) and the host gene expression in control conditions. With a Mann-Whitney U test we checked that the properties of the windows previously described were balanced between the two sets.

CircRNAs with at least 2 overlapping TDP-binding regions in 2 independent datasets were defined as TDP interactors.

Enrichment of TDP-43 binding within c-circRNA flanking introns relative to matched control circRNAs was assessed using Fisher’s exact test.

### Cryptic Exon mapping analysis

Genomic coordinates of TDP-43-dependent cryptic splicing events were obtained from Brown et al.^15^, including both exon inclusion and exon skipping events. Coordinates of the c-circRNAs flanking introns were extracted from the corresponding transcript annotations. The same set of matched control circRNAs used for the TDP-43 binding analysis was employed as background. A cryptic event was considered associated with a circRNA when its genomic coordinates overlapped, in a strand-specific manner, either of the two flanking introns. Enrichment of cryptic splicing events within c-circRNA flanking introns relative to matched control circRNAs was assessed using Fisher’s exact test.

### NYGC ALS Consortium RNA-sequencing cohort

For our post-mortem tissue RNA-sequencing we used samples from the NYGC ALS Consortium, including non-neurological disease controls, FTLD, ALS, FTD with ALS (ALS-FTLD), or ALS with suspected Alzheimer’s disease (ALS-AD). Patients with FTD were classified according to a pathologist’s diagnosis of FTD with TDP-43 inclusions (FTLD-TDP), or those with FUS or Tau aggregates. ALS samples were divided into the following subcategories using the available Consortium metadata: ALS with or without reported SOD1 or FUS mutations. All non-SOD1 or FUS ALS samples were grouped as ALS-TDP in this work for simplicity, although reporting of postmortem TDP-43 inclusions was not systematic and therefore not integrated into the metadata. Confirmed TDP-43 pathology postmortem was reported for all FTLD-TDP samples.

Sample processing, library preparation, and RNA-seq quality control have been extensively described in previous papers^24,30,59^. In brief, RNA was extracted from flash-frozen postmortem tissue using TRIzol (Thermo Fisher Scientific) chloroform, and RNA-Seq libraries were prepared from 500 ng total RNA using the KAPA Stranded RNA-Seq Kit with RiboErase (KAPA Biosystems) for ribosomal RNA depletion. Pooled libraries (average insert size: 375 bp) passing the quality criteria were sequenced either on an Illumina HiSeq 2500 (125 bp paired end) or an Illumina NovaSeq (100 bp paired end). The samples had a median sequencing depth of 42 million read pairs, with a range between 16 and 167 million read pairs.

Samples were uniformly processed, including adapter trimming with Trimmomatic and alignment to the hg38 genome build using STAR (2.7.2a) with indexes from GENCODE v30. Extensive quality control was performed using SAMtools and Picard Tools to confirm sex and tissue of origin.

Potential confounding variables were assessed independently for each tissue. Differences in sequencing depth, RNA integrity number (RIN), and age at death between groups were evaluated using two-sided Wilcoxon–Mann–Whitney tests followed by Bonferroni correction for multiple testing. Sex distribution was compared using Fisher’s exact test.

### NYGC circRNA detection and enrichment analysis

CircRNAs were detected as described in the section “CircRNA detection”. A circRNA was considered expressed in a sample if at least one read supporting the back-splicing junction (BSJ) was detected. To identify circRNAs enriched in ALS-TDP samples, we compared the proportion of samples expressing each circRNA between ALS-TDP and control groups independently for each tissue using Fisher’s exact test. P-values were corrected for multiple testing using the Benjamini–Hochberg procedure. CircRNAs enriched in ALS-TDP samples and detected in fewer than 5% of control samples were classified as cryptic circRNAs (c-circRNAs).

For individual c-circRNA candidates, expression was quantified as reads per million mapped reads (RPM), calculated from BSJ-supporting reads normalized to the total number of mapped reads. Corresponding linear transcripts were quantified using Salmon (v1.8.0) and expressed as transcripts per million (TPM). Differences between groups were assessed independently for each tissue using two-sided Wilcoxon–Mann–Whitney tests followed by Bonferroni correction for multiple testing.

Heatmaps were generated using the ComplexHeatmap^58^ R package (v2.8.0).

### TDP-43 cryptic burden analysis

TDP-43 cryptic burden scores were obtained as previously described^15^. For each tissue, samples were ranked according to their burden score and divided into a low-burden group (bottom 25%) and a high-burden group (top 25%). Differences in sequencing depth between burden groups were assessed independently for each tissue using two-sided Wilcoxon– Mann–Whitney tests followed by Bonferroni correction. Enrichment of circRNAs between high- and low-burden groups was evaluated independently for each tissue using Fisher’s exact test by comparing the proportion of samples expressing each circRNA between the two groups.

### Statistical software and visualization

All statistical analyses were performed using R^60^ (v4.5.1). Receiver operating characteristic (ROC) curve analyses were conducted using pROC^61^ (v1.19.0.1). Graphical representations were generated using the R packages ggplot2^62^ (v4.0.1), ComplexHeatmap^58^ (v2.24.1), EnhancedVolcano^63^ (v1.26.0), eulerr^64^ (v7.0.4), circlize^65^ (v0.4.16), ggvenn^66^ (v0.1.19), ggpubr^67^ (v0.6.2), RColorBrewer^68^ (v1.1.3), and viridisLite^69^ (v0.4.2). The distribution and deviation of data shown in the figures of this work, the statistical tests used to calculate significant differences, and the exact value of n (e.g., the number of biological replicates of the experiments) are denoted in figure legends. In figure legends “SD” stands for “standard deviation” and “SEM” stands for “standard error mean”. Significance values were depicted in the figures using the following key legend: n.s. p>0.05, *p < 0.05, **p < 0.01, ***p < 0.001. In the box plots interquartile range spans from 75^th^ percentile and 25th percentile of data with the median indicated as line in the box (50th percentile).

When needed, data were plotted using GraphPad Prism 10.

## Data availability

Publicly available data were obtained from ArrayExpress under the accession numbers E-MTAB-15433 (i3LMN with TDP-43 knockdown) and from the Sequence Read Archive (SRA) under the bioproject PRJNA1256902 (SH-SY5Y cells with TDP-43 knockdown and nonsense-mediated decay inhibition). RNA-seq data generated by the NYGC ALS Consortium and used in this study can be accessed through the GEO (GSE137810, GSE124439, GSE116622 and GSE153960).

## Acknowledgements

This work was supported by Target ALS (PF; MEW); National Institute of Health U54NS123743 (PF); Packard Center for ALS Research (PF); Trace Neuroscience Inc (PF).

