## Supplementary Figures for "TDP-43 dysfunction induces cryptic circular RNAs in ALS/FTD"

**A**

| | Description | TDP-KD Method | n CTRL samples | n KD samples |
|---|---|---|---|---|
| **SH-SY5Y** | Neuroblastoma cell line | siRNA | 3 | 3 |
| **SK-N-BE(2)** | Neuroblastoma cell line | siRNA | 3 | 3 |
| **i3LMN** | hiPSCs-derived lower motor nuerons | CRISPRi | 8 | 8 |
| **i3N** | hiPSCs-derived cortical nuerons | CRISPRi | 11 | 12 |

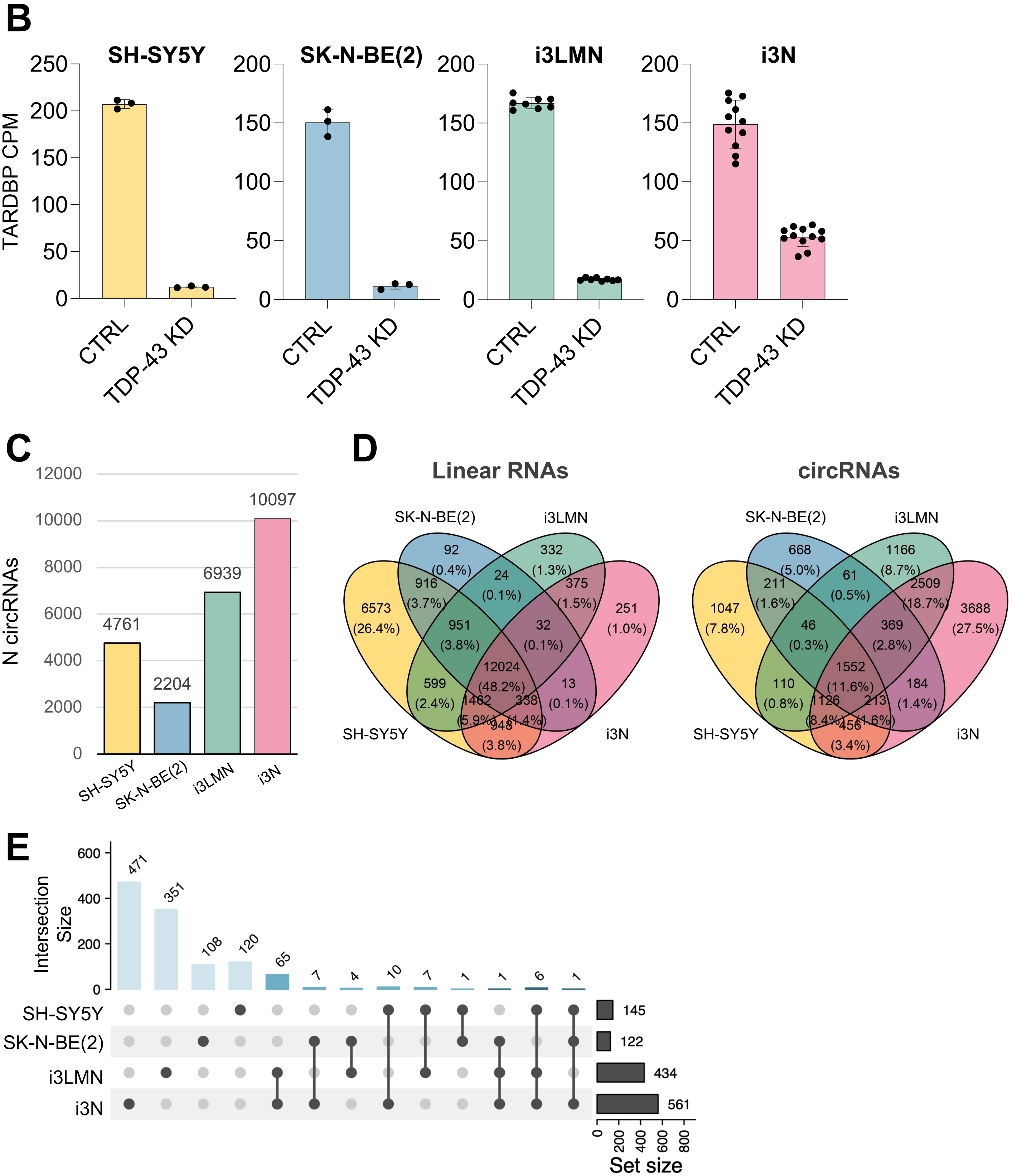

**Figure S1: A)** Table describing the RNA-seq datasets used in this study, indicating the cellular model, TDP-43 knockdown strategy and the number of control (CTRL) and TDP-43 knockdown (KD) samples analysed. **B)** RNA-seq quantification of TARDBP expression across the four datasets under control (CTRL) and TDP-43 knockdown (TDP-43 KD) conditions. Bars represent the mean counts per million (CPM), with individual biological replicates shown as black dots. Error bars indicate ± SD. **C)** Bar plot showing the number of circRNAs detected by CIRI2 in each dataset. **D)** Venn diagrams showing the overlap of linear RNAs (left) and circRNAs (right) detected across the four cellular models. Numbers indicate transcripts unique to or shared among the different datasets, with percentages reported relative to the total number of detected linear RNAs or circRNAs, respectively. Compared with linear RNAs, circRNAs exhibited substantially greater cell-type specificity, with fewer than 12% detected in all four models. **E)** UpSet plot showing the overlap of downregulated circRNAs identified across four cellular models: SH-SY5Y, SK-N-BE(2), i3LMNs, and i3N. Horizontal bars on the right indicate the total number of downregulated circRNAs detected in each model, whereas vertical bars represent the size of each intersection. Connected dots denote the datasets contributing to each intersection.

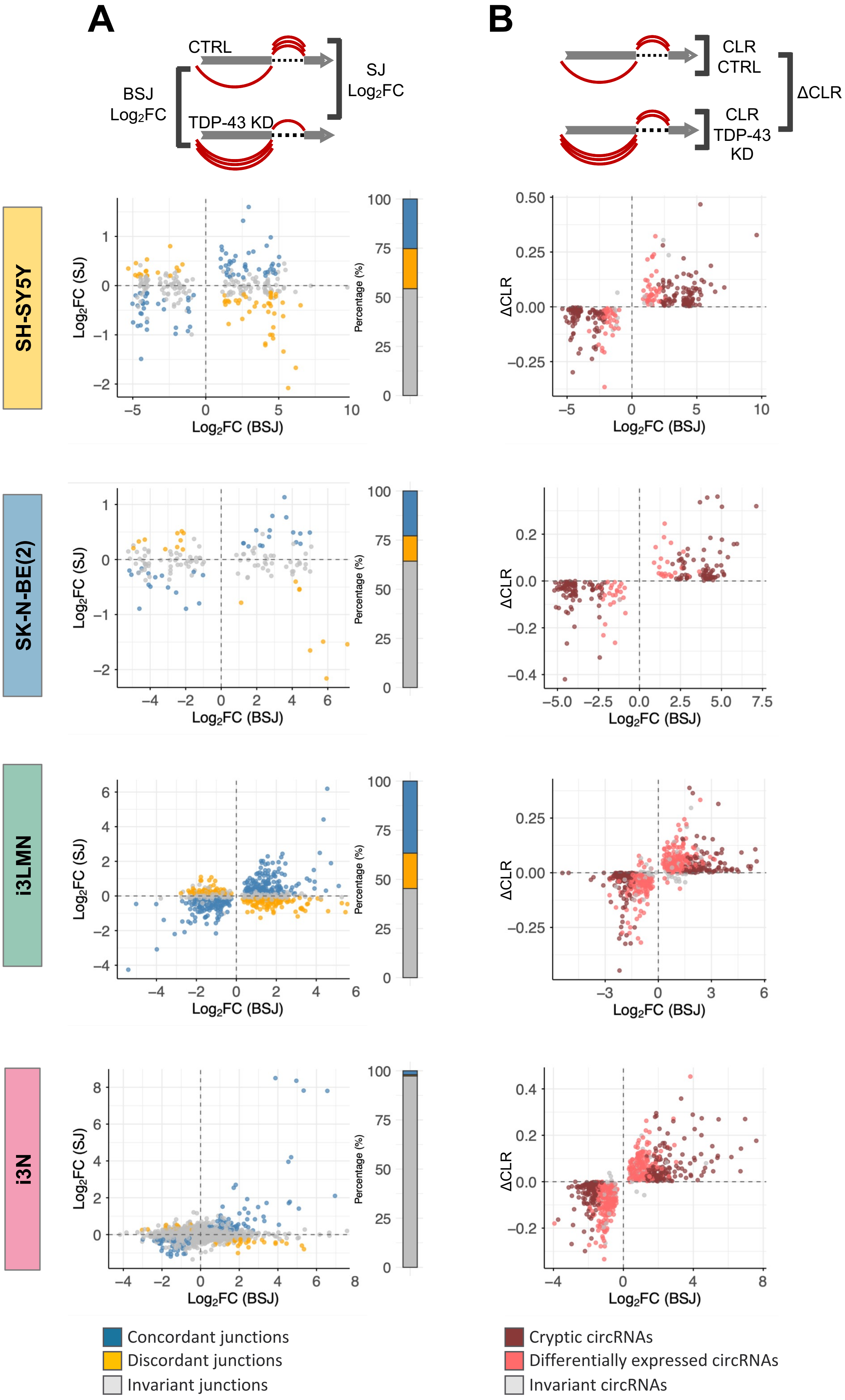
A
B
CTRL
TDP-43 KD
BSJ
Log2FC
SJ
Log2FC
CLR
CTRL
CLR
TDP-43
KD
ΔCLR
SH-SY5Y
SK-N-BE(2)
i3LMN
i3N
Log2FC (SJ)
Log2FC (BSJ)
Percentage (%)
ΔCLR
Concordant junctions
Discordant junctions
Invariant junctions
Cryptic circRNAs
Differentially expressed circRNAs
Invariant circRNAs

**Figure S2: A)** Left panel: scatter plot comparing the $\log_2$ fold change ($\log_2$FC) of circRNAs (back-splice junctions, BSJs; x-axis) and their cognate linear splice junctions (SJs; y-axis) following TDP-43 knockdown. CircRNAs were classified as invariant when the corresponding linear isoform showed no significant change (grey), concordant when both circular and linear transcripts changed in the same direction (blue), and discordant when they changed in opposite directions (yellow). Right panel: bar plot summarising the proportion of invariant, concordant and discordant circRNAs in each cellular model. **B)** Scatter plot comparing circRNA differential expression ($\log_2$FC(BSJ); x-axis) with the corresponding change in circular-to-linear ratio ($\Delta$CLR; y-axis) upon TDP-43 depletion. Invariant circRNAs are shown in grey, differentially expressed circRNAs in red, and cryptic circRNAs in dark red.

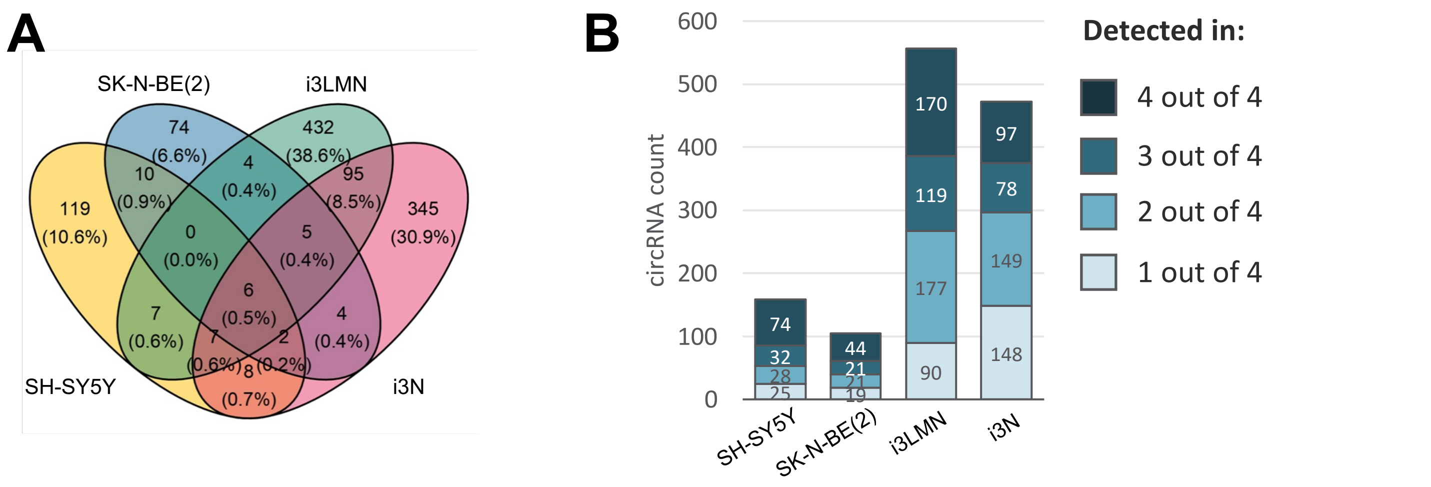


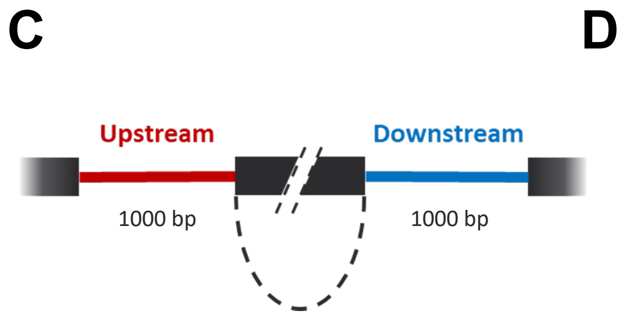


D

| Name | Experiment | Model | Peak_Caller |
|---|---|---|---|
| CLIP1 | iCLIP | SH-SY5Y | Piranha |
| CLIP2 | iCLIP | SH-SY5Y | CIMS |
| CLIP3 | iCLIP | Brain | Piranha |
| CLIP4 | iCLIP | Brain | CIMS |
| CLIP5 | iCLIP | H9 | CIMS |
| CLIP6 | iCLIP | H9 | Piranha |
| CLIP7 | PAR CLIP | HEK293T | PARalyzer |
| CLIP8 | PAR CLIP | HEK293T | Piranha |
| CLIP9 | eCLIP | K562 | |

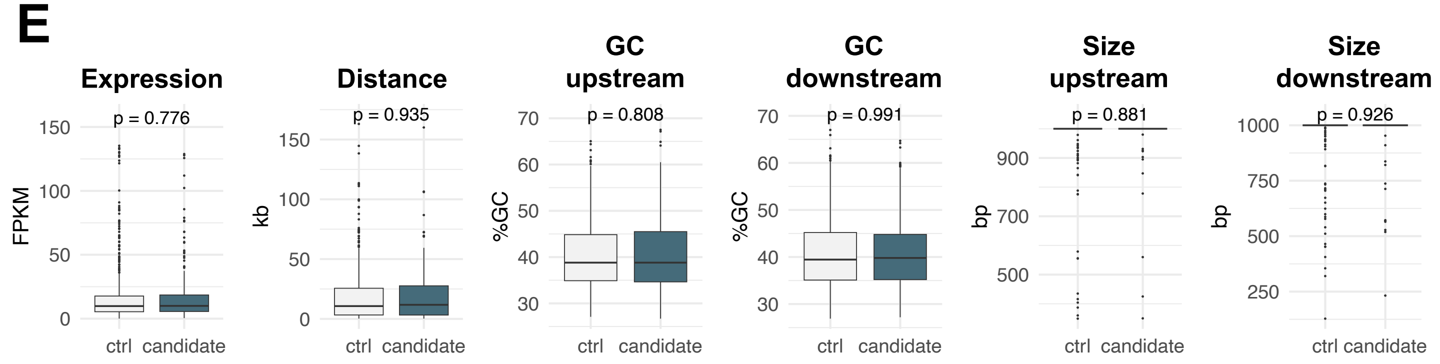


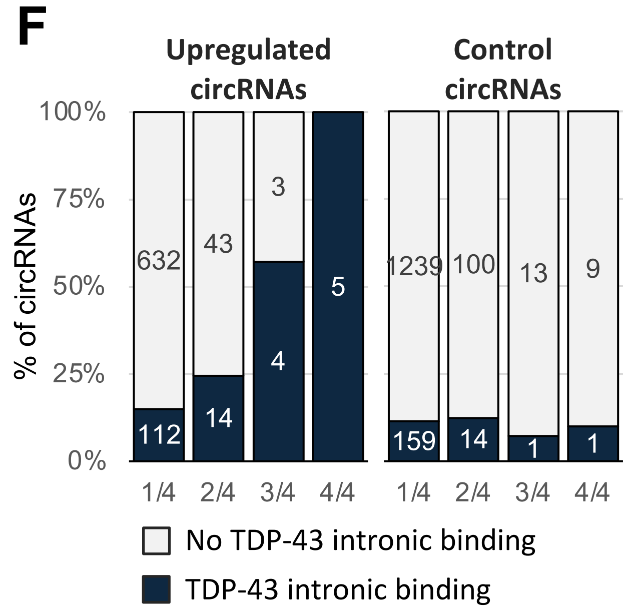


Fisher's Exact Test

| | OR | pvalue |
|---|---|---|
| 1/4 | 1.38 | 0.017 |
| 2/4 | 2.31 | 0.050 |
| 3/4 | 14.39 | 0.025 |
| 4/4 | Inf | 0.002 |

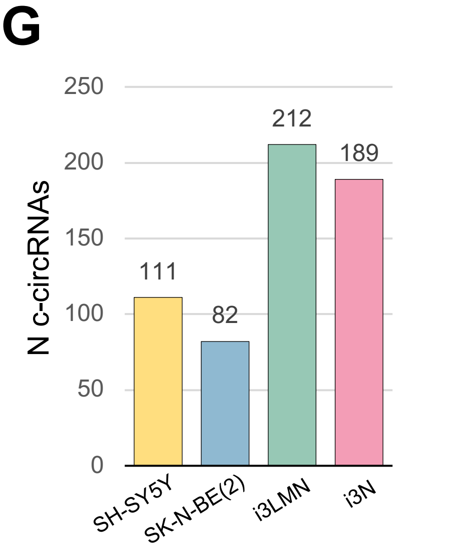

**Figure S3: A)** Venn diagrams showing the overlap of cryptic circRNAs (c-circRNAs) detected across the four cellular models: SH-SY5Y, SK-N-BE(2), i3LMNs and i3Ns. Numbers indicate c-circRNAs unique to or shared among the different models, with percentages reported relative to the total number of c-circRNAs detected. **B)** Stacked bar plot showing the distribution of upregulated circRNAs according to the number of cellular models in which they were detected, ranging from cell-type-specific (1/4) to shared across all four models (4/4). Numbers in the bars indicate the corresponding circRNA counts. **C)** Schematic representation of the strategy used to assess TDP-43 binding within the introns flanking candidate circRNAs. A region of 1000 bp either upstream or downstream the back-splice junction was interrogated for TDP-43 CLIP peaks. **D)** Table summarising the POSTAR3 CLIP datasets used to retrieve TDP-43 binding peaks. **E)** Characterisation of the candidate (upregulated) and control (invariant) circRNA sets used for TDP-43 binding enrichment analysis. Boxplots compare host gene expression under control conditions (FPKM), the distance between circularising exons (kb), and the GC content (%GC) and length (bp) of the upstream and downstream flanking introns. No significant differences were observed between the two groups (two-sided Wilcoxon rank-sum test; p values shown), confirming that the matched control circRNAs provide an appropriate background for TDP-43 binding enrichment analysis. **F)** Stacked bar plot showing the proportion of upregulated circRNAs following TDP-43 knockdown that either exhibit (blue) or lack (grey) TDP-43 binding within their flanking intron. CircRNAs are grouped according to the number of datasets in which they were detected. For each group, TDP-43 binding is compared with a control set of invariant circRNAs (two for every candidate circRNA). Numbers in the bars indicate the corresponding circRNA counts. Odds ratios and P values were calculated using Fisher's exact test. **G)** Bar plot showing the number of cryptic circRNAs identified in each cellular model.

A

| | Control | ALS-non-TDP | ALS-TDP | FTD-non-TDP | FTD-TDP | TOT |
|---|---|---|---|---|---|---|
| Cervical Spinal Cord | 56 | 12 | 331 | 0 | 0 | 399 |
| Lumbar Spinal Cord | 54 | 12 | 310 | 0 | 0 | 376 |
| Motor Cortex | 71 | 18 | 445 | 0 | 0 | 534 |
| Frontal Cortex | 84 | 11 | 306 | 11 | 37 | 449 |
| Temporal Cortex | 27 | 2 | 35 | 8 | 33 | 105 |

**TOTAL = 1863 samples**

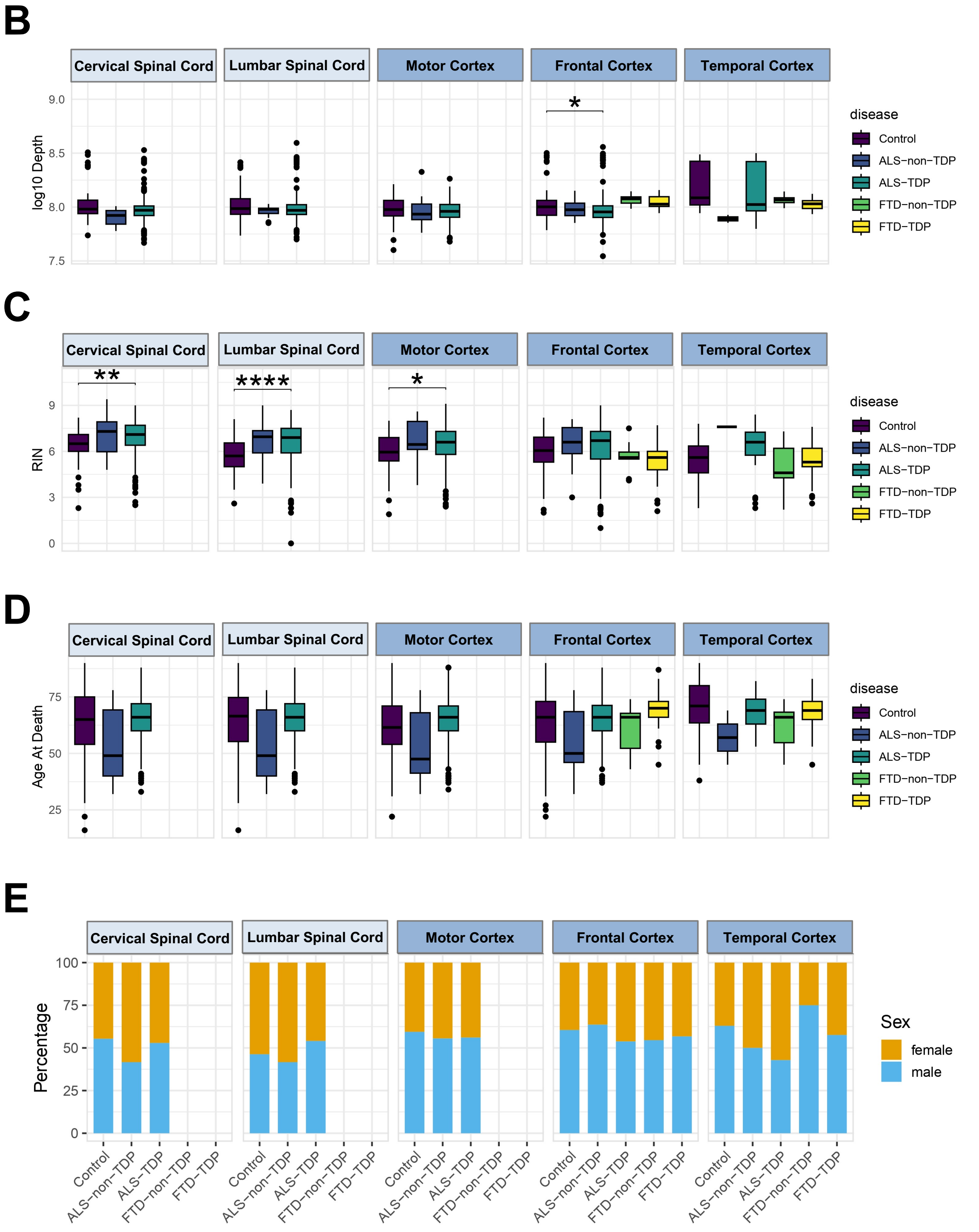

**Figure S4: A)** Summary of the NYGC ALS Consortium cohort analysed in this study. The table shows the number of post-mortem tissue samples from healthy controls, ALS and FTD cases with or without TDP-43 pathology across the CNS regions analysed. The total number of samples for each tissue is indicated. **B–D)** Box plots comparing technical and demographic variables across the analysed samples. Total RNA-seq library size (log10-transformed; **B**), RNA integrity number (RIN; **C**), and age at death (**D**) are shown. Statistical significance was assessed using the Wilcoxon-Mann-Whitney test with Bonferroni correction (*$p < 0.05$). Non-significant comparisons ($p > 0.05$) are not reported. **E)** Stacked bar plots showing the sex distribution across disease groups for each CNS region analysed. Statistical significance was assessed using Fisher's exact test. Non-significant comparisons ($p > 0.05$) are not reported.

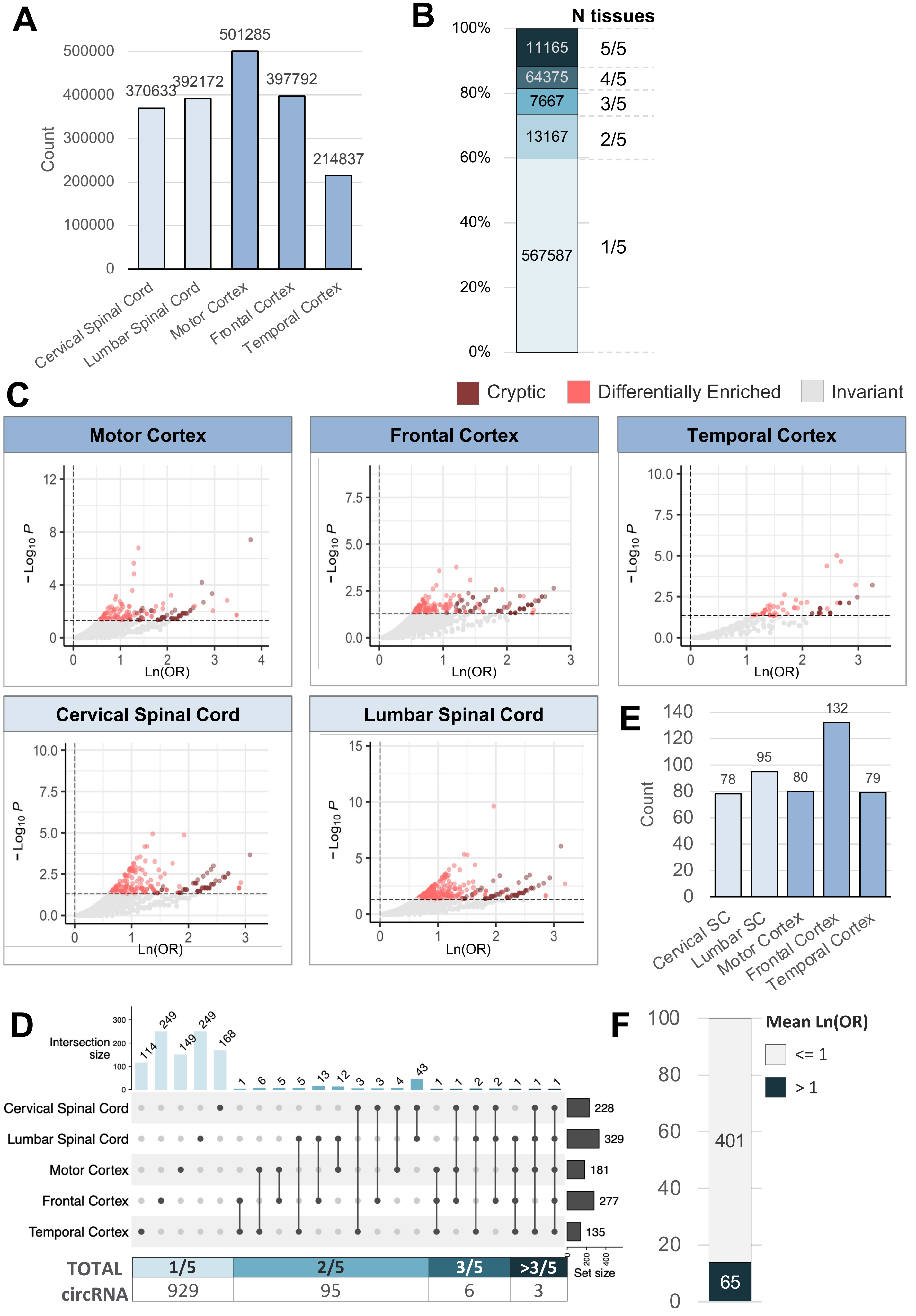
A
Count
370633
392172
501285
397792
214837
Cervical Spinal Cord
Lumbar Spinal Cord
Motor Cortex
Frontal Cortex
Temporal Cortex
B
N tissues
11165
64375
7667
13167
567587
5/5
4/5
3/5
2/5
1/5
C
Cryptic
Differentially Enriched
Invariant
Motor Cortex
Frontal Cortex
Temporal Cortex
Cervical Spinal Cord
Lumbar Spinal Cord
Ln(OR)
D
Intersection size
Set size
228
329
181
277
135
TOTAL circRNA
1/5
2/5
3/5
>3/5
929
95
6
3
E
Count
78
95
80
132
79
Cervical SC
Lumbar SC
Motor Cortex
Frontal Cortex
Temporal Cortex
F
Mean Ln(OR)
<= 1
> 1
401
65

**Figure S5: A)** Bar plot showing the number of circRNAs detected in each spinal cord (light blue) and cortical (dark blue) tissue analysed. **B)** Stacked bar plot showing the distribution of circRNAs according to the number of CNS regions in which they were detected, ranging from tissue-specific (1/5) to shared across all five tissues (5/5). Numbers indicate the corresponding circRNA counts. **C)** Volcano plots showing the enrichment of circRNAs in ALS-TDP samples relative to controls across the five CNS regions analysed. The x-axis represents the natural logarithm of the odds ratio (lnOR), and the y-axis the −log10 P value from Fisher's exact test. Invariant circRNAs are shown in grey, differentially enriched circRNAs in red, and cryptic circRNAs in dark red. Horizontal and vertical dashed lines indicate the significance and enrichment thresholds, respectively. **D)** UpSet plot showing the overlap of differentially enriched circRNAs detected across the five CNS regions. Horizontal bars on the right indicate the total number of differentially enriched circRNAs identified in each tissue, whereas vertical bars represent the size of each intersection. Connected dots denote the tissues contributing to each intersection. The lower panel shows the number of differentially enriched circRNAs identified in one (1/5), two (2/5), three (3/5), or more than three (>3/5) tissues. **E)** Bar plot showing the number of cryptic circRNAs identified in each spinal cord (light blue) and cortical (dark blue) tissue. **F)** Stacked bar plot showing the distribution of cryptic circRNAs according to their mean enrichment across tissues. Blue indicates c-circRNAs with a mean natural logarithm of the odds ratio (lnOR) > 1, whereas grey indicates c-circRNAs with a mean lnOR ≤ 1.

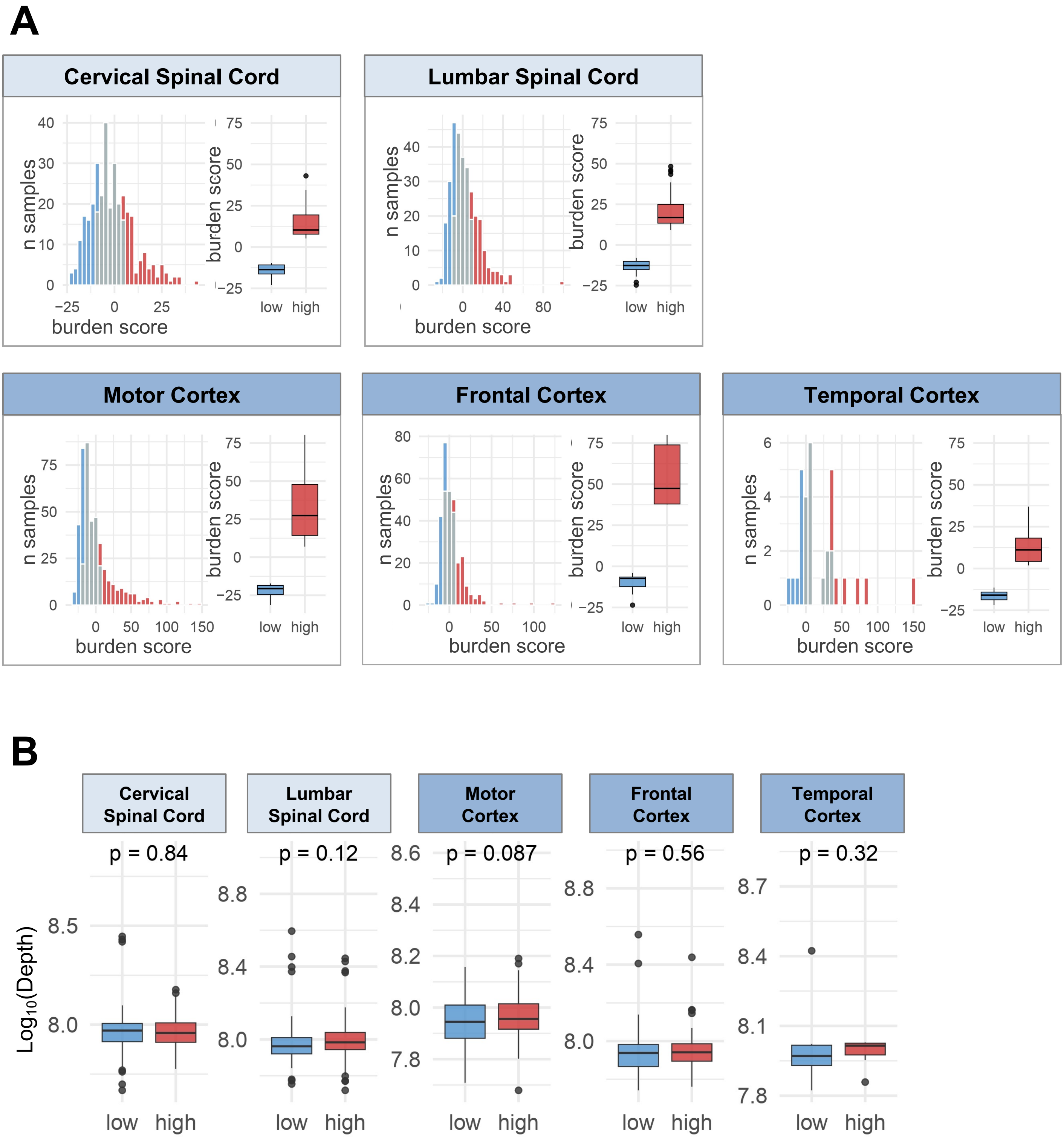
A
Cervical Spinal Cord
Lumbar Spinal Cord
Motor Cortex
Frontal Cortex
Temporal Cortex
n samples
burden score
low
high
B
Cervical Spinal Cord
Lumbar Spinal Cord
Motor Cortex
Frontal Cortex
Temporal Cortex
p = 0.84
p = 0.12
p = 0.087
p = 0.56
p = 0.32
Log10(Depth)
low
high

**Figure S6: A)** Histograms showing the distribution of TDP-43 cryptic burden scores across ALS-TDP samples for each CNS region analysed. Samples were stratified into “low” (first quartile), “medium” (second and third quartiles), and “high” (fourth quartile) burden groups. Box plots show the distribution of burden scores in the low- and high-burden groups used for subsequent analyses. **B)** Box plots comparing total RNA-seq library size (log10-transformed) between the low- and high-burden groups across the five CNS regions. Statistical significance was assessed using the Wilcoxon–Mann–Whitney test with Bonferroni correction.

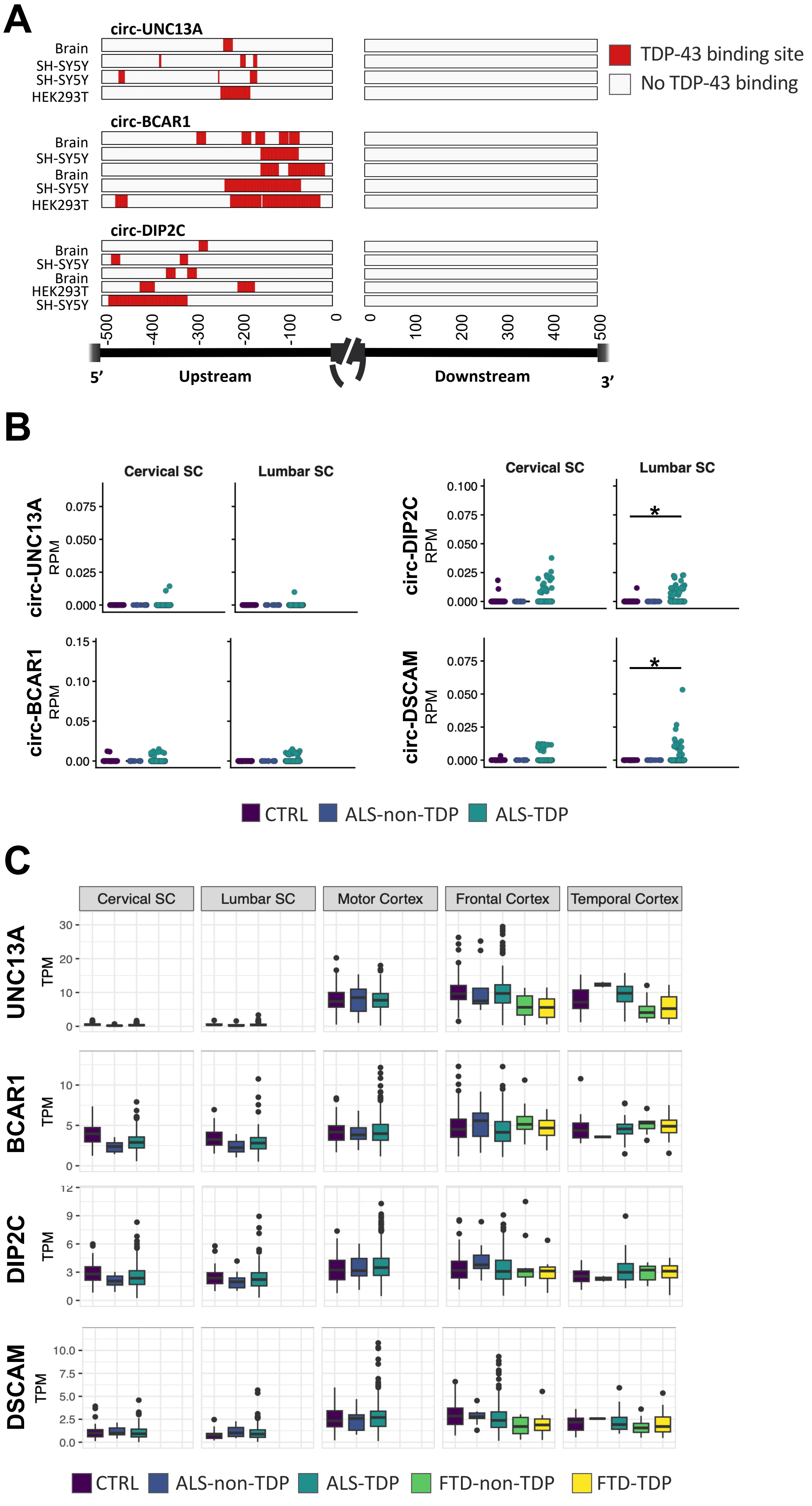

A
circ-UNC13A
Brain
SH-SY5Y
SH-SY5Y
HEK293T
circ-BCAR1
Brain
SH-SY5Y
Brain
SH-SY5Y
HEK293T
circ-DIP2C
Brain
SH-SY5Y
Brain
HEK293T
SH-SY5Y
TDP-43 binding site
No TDP-43 binding
5'
Upstream
Downstream
3'
B
Cervical SC
Lumbar SC
circ-UNC13A RPM
circ-BCAR1 RPM
circ-DIP2C RPM
circ-DSCAM RPM
CTRL
ALS-non-TDP
ALS-TDP
C
Cervical SC
Lumbar SC
Motor Cortex
Frontal Cortex
Temporal Cortex
UNC13A TPM
BCAR1 TPM
DIP2C TPM
DSCAM TPM
CTRL
ALS-non-TDP
ALS-TDP
FTD-non-TDP
FTD-TDP

**Figure S7: A)** Distribution of TDP-43 binding sites within the flanking introns of selected c-circRNAs (c-circUNC13A, c-circBCAR1, c-circDIP2C), based on CLIP-seq datasets from indicated cells and tissues retrieved from POSTAR3. Red regions indicate experimentally identified TDP-43 binding sites, whereas grey regions indicate regions without detected TDP-43 binding. Coordinates are shown relative to the back-splice junction. **B)** Box plots showing the expression of the selected c-circRNAs (c-circUNC13A, c-circBCAR1, c-circDIP2C, and c-circDSCAM) in ALS-TDP, ALS-non-TDP and control spinal cord samples from the NYGC ALS Consortium. Expression of the c-circRNAs was quantified from back-splice junction (BSJ) counts normalised for library size (reads per million, RPM). Each dot represents an individual sample. Statistical significance was assessed using Wilcoxon-Mann-Whitney test with Bonferroni correction (*p < 0.05, **p < 0.01, ***p < 0.001). Non-significant comparisons (p > 0.05) are not reported. **C)** Box plots showing the expression of the linear host transcripts corresponding to the candidate c-circRNAs (UNC13A, BCAR1, DIP2C, and DSCAM) in the NYGC ALS Consortium RNA-seq cohort across disease groups and CNS regions. Gene expression was quantified as transcripts per million (TPM). Each dot represents an individual sample. Statistical significance was assessed using the Wilcoxon–Mann–Whitney test with Bonferroni correction (*p < 0.05, **p < 0.01, ***p < 0.001). Non-significant comparisons are not shown.

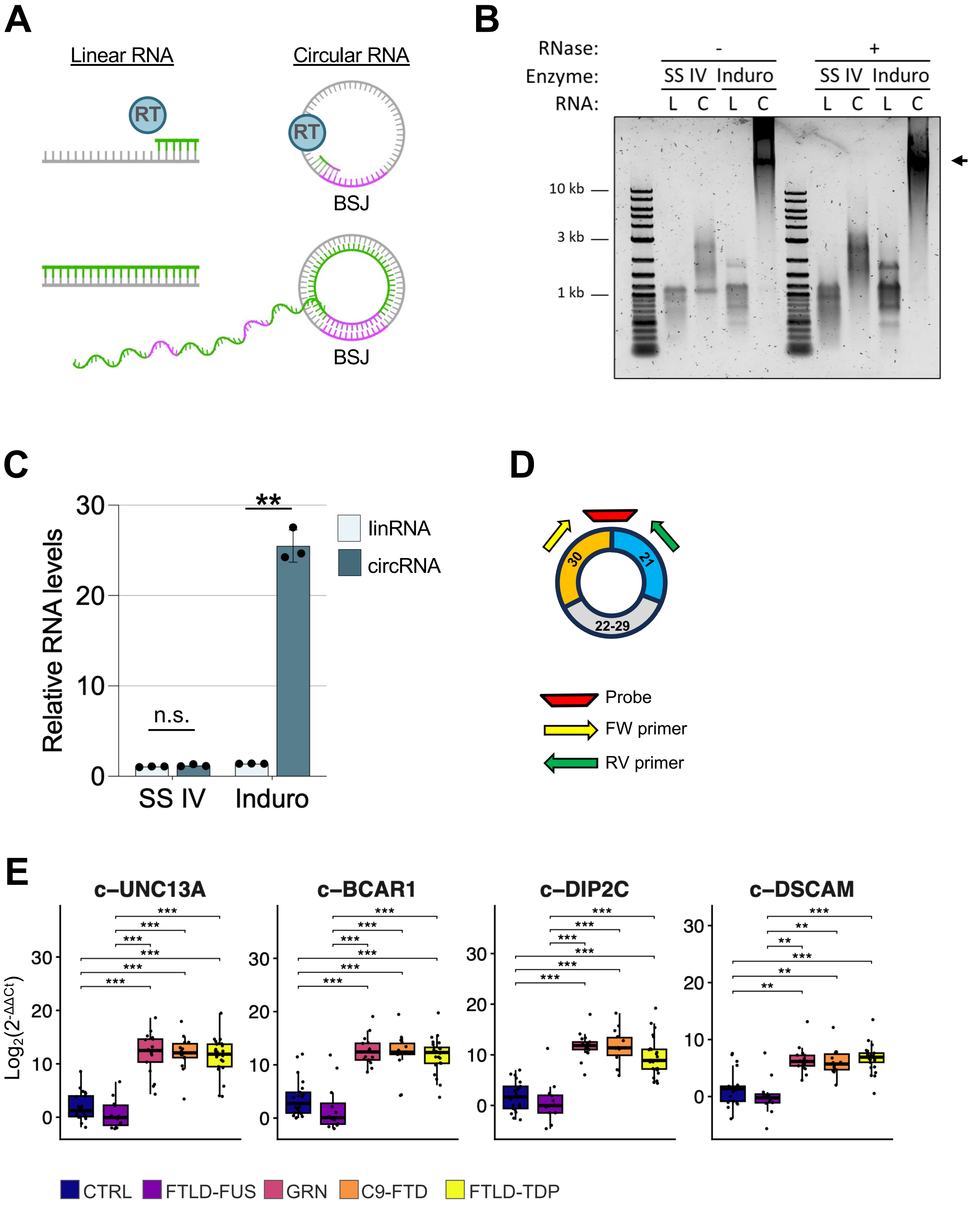

A
Linear RNA
Circular RNA
RT
BSJ
B
RNase: - +
Enzyme: SS IV Induro SS IV Induro
RNA: L C L C L C L C
10 kb
3 kb
1 kb
C
Relative RNA levels
30
20
10
0
**
n.s.
SS IV
Induro
linRNA
circRNA
D
30
21
22-29
Probe
FW primer
RV primer
E
c–UNC13A
c–BCAR1
c–DIP2C
c–DSCAM
$Log_2(2^{-\Delta\Delta Ct})$
***
**
CTRL
FTLD-FUS
GRN
C9-FTD
FTLD-TDP

**Figure S8: A)** Schematic illustrating reverse transcription of linear and circular RNA templates by the Group II intron reverse transcriptase Induro. Rolling-circle reverse transcription of circular RNAs generates long cDNA molecules containing multiple copies of the back-splice junction (BSJ). **B)** Agarose gel showing cDNA products generated by reverse transcription of in vitro-transcribed linear or circular UNC13A RNA using either SuperScript IV (SS IV), which does not support rolling-circle reverse transcription, or Induro. Samples were analysed before and after RNase H/RNase I treatment. Long concatemeric cDNAs, indicated by the black arrow, generated by rolling-circle reverse transcription are observed exclusively following reverse transcription of the circular RNA with Induro. **C)** RT–qPCR analysis of in vitro transcribed linear ("linRNA", light blue) or circular ("circRNA", dark blue) UNC13A RNA following reverse transcription with SuperScript IV (SS IV) or Induro. Expression is presented as relative values, with the SS IV linear RNA condition set to 1. Bars represent the mean ± SD of three independent experiments ($n = 3$). Statistical significance was assessed using a two-sided Student's t-test (n.s. $p > 0.05$; **$p < 0.01$). **D)** Schematic representation of the RT-qPCR detection strategy used to quantify cryptic circRNAs (c-circRNAs) in post-mortem human tissues, illustrated using c-circUNC13A as an example. Divergent primers specifically amplify the circular transcript, while a TaqMan probe spanning the back-splice junction (BSJ) ensures selective detection. **E)** RT–qPCR analysis of the selected c-circRNAs (c-circUNC13A, c-circBCAR1, c-circDIP2C, c-circDSCAM) in post-mortem human frontal cortex samples from healthy controls (n=21) and FTD patients with (GRN n=17, C9-FTD n=14, FTLD-TDP n=25) or without TDP-43 pathology (FTD-FUS n=12). c-circRNA expression was normalised to GAPDH mRNA and is presented as fold change ($\log_2(2^{-\Delta\Delta Ct})$), with respect to a control sample set to a value of 1. Each dot represents an individual sample. Statistical significance was assessed using Wilcoxon-Mann-Whitney test with Bonferroni correction (*$p < 0.05$, **$p < 0.01$, ***$p < 0.001$). Non-significant comparisons ($p > 0.05$) are not shown.

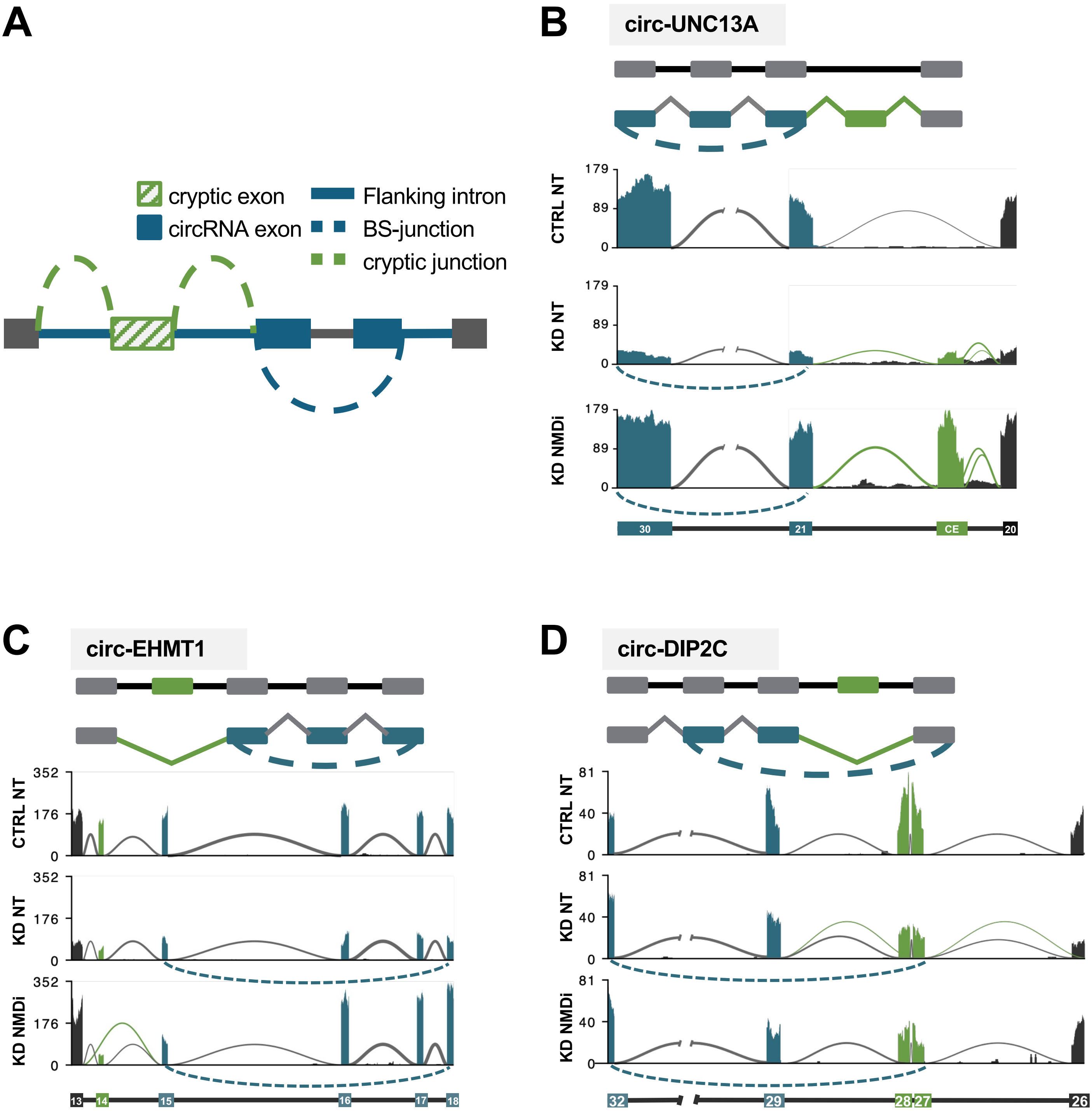

A
cryptic exon
circRNA exon
Flanking intron
BS-junction
cryptic junction
B
circ-UNC13A
CTRL NT
KD NT
KD NMDi
179
89
0
30
21
CE
20
C
circ-EHMT1
352
176
0
13
14
15
16
17
18
D
circ-DIP2C
81
40
0
32
29
28
27
26

**Figure S9: A)** Schematic representation of the meta-circRNA used to assess the presence of cryptic splice junctions within the introns flanking High-Confidence cryptic circRNAs (c-circRNAs) identified in Fig. 3A. The positions of the back-splice junction (BSJ), cryptic splice junction, circularising exon(s), and flanking introns are indicated. **B-D)** Sashimi plot showing the concomitant occurrence of a cryptic splicing event (green) and a cryptic circular RNA (c-circRNA)–forming event (blue) in the UNC13A (B), EHMT1 (C) and DIP2C (D) genes. The upper panel shows a schematic representation of the events, while the lower panels display read coverage and splice junction usage under the indicated conditions: control (CTRL NT), TDP-43 knockdown (KD NT), and TDP-43 knockdown with nonsense-mediated decay inhibition (KD NMDi).

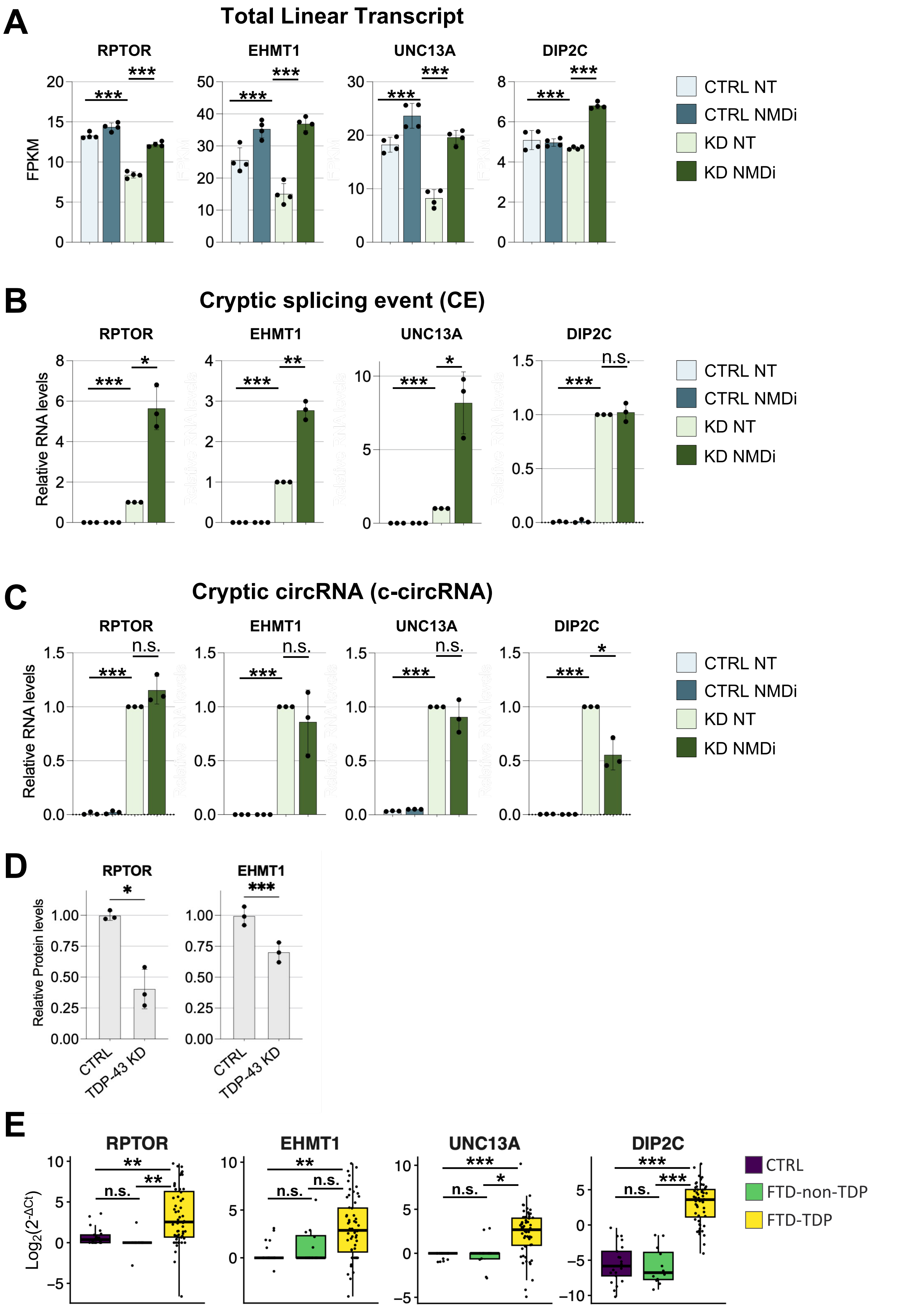
A
Total Linear Transcript
RPTOR
EHMT1
UNC13A
DIP2C
FPKM
***
CTRL NT
CTRL NMDi
KD NT
KD NMDi
B
Cryptic splicing event (CE)
RPTOR
EHMT1
UNC13A
DIP2C
Relative RNA levels
n.s.
C
Cryptic circRNA (c-circRNA)
RPTOR
EHMT1
UNC13A
DIP2C
Relative RNA levels
n.s.
D
RPTOR
EHMT1
Relative Protein levels
CTRL
TDP-43 KD
E
RPTOR
EHMT1
UNC13A
DIP2C
$Log_2(2^{-\Delta Ct})$
n.s.
CTRL
FTD-non-TDP
FTD-TDP

**Figure S10: A)** Expression of total linear transcripts of RPTOR, EHMT1, UNC13A and DIP2C, quantified as fragments per kilobase of transcript per million mapped reads (FPKM), from bulk RNA-seq data generated from SH-SY5Y cells under control (CTRL) or TDP-43 knockdown (KD) conditions, with (NMDi) or without (NT) inhibition of nonsense-mediated decay (NMD) by cycloheximide (CHX). Bars represent the mean ± SD of independent biological replicates, with individual replicates shown as black dots (n=4). Statistical significance was assessed using two-tailed unpaired Student's *t* test with correction for multiple test comparison (FDR Benjamini–Hochberg) (***p < 0.001). **B,C)** RT–qPCR analysis of cryptic linear RNAs (CE; B) and their corresponding cryptic circRNAs (c-circRNA; C) derived from RPTOR, EHMT1, UNC13A and DIP2C. RNA was extracted from SH-SY5Y cells under control (CTRL) or TDP-43 knockdown (KD) conditions, with (NMDi) or without (NT) inhibition of nonsense-mediated decay (NMD) using SMG-1 inhibitor 11j. Transcript abundance was normalised to GAPDH mRNA and is presented as relative expression values (2^−ΔΔCt), with respect to the KD NT set to a value of 1. Each dot represents an independent biological replicate (n= 3. Statistical significance was assessed using a two-tailed unpaired Student's *t* test with correction for multiple test comparison (FDR Benjamini–Hochberg) (n.s. p > 0.05, *p < 0.05, **p < 0.01,***p < 0.001). **D)** Quantification of RPTOR and EHMT1 protein levels from three independent biological replicates (n=3), one of which is shown as a representative immunoblot in Fig. 3D. Protein levels were normalised to α-Tubulin and are presented with respect to a control sample set to a value of 1. Statistical significance was assessed using a two-tailed unpaired Student's *t*-test (*p < 0.05, ***p < 0.001). **E)** c-circRNA:CE ratios calculated from the RT–qPCR data shown in Fig. 3F for the indicated c-circRNAs and their corresponding CEs in post-mortem human frontal cortex samples from healthy controls (CTRL, *n* = 21), FTD patients without TDP-43 pathology (FTD-non-TDP, *n* = 12), and FTD patients with TDP-43 pathology (FTD-TDP, *n* = 56). Ratios are presented as $\log_2$(2^−ΔCt), where ΔCt = Ct(c-circRNA) − Ct(CE) within the same sample. Each dot represents an individual sample. Statistical significance was assessed using Wilcoxon-Mann-Whitney test with Bonferroni correction (n.s. p > 0.05, *p < 0.05, **p < 0.01,***p < 0.001).

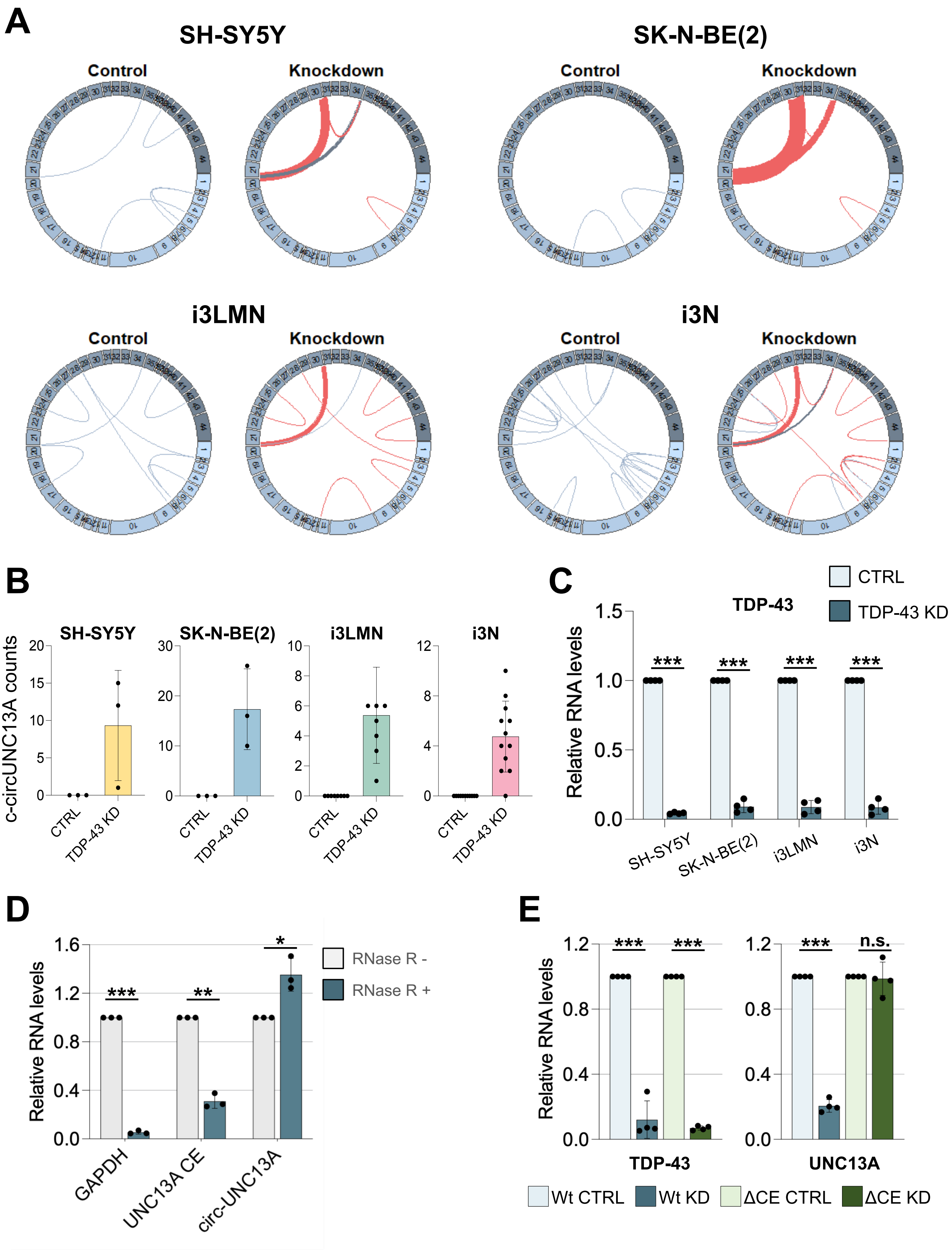

A
SH-SY5Y
SK-N-BE(2)
Control
Knockdown
i3LMN
i3N
B
c-circUNC13A counts
CTRL
TDP-43 KD
C
TDP-43
CTRL
TDP-43 KD
Relative RNA levels
***
SH-SY5Y
SK-N-BE(2)
i3LMN
i3N
D
Relative RNA levels
RNase R -
RNase R +
GAPDH
UNC13A CE
circ-UNC13A
***
**
*
E
Relative RNA levels
TDP-43
UNC13A
n.s.
Wt CTRL
Wt KD
ΔCE CTRL
ΔCE KD

**Figure S11: A)** Schematic representation of UNC13A circRNA isoforms detected across the four cellular models under control (Control) and TDP-43 knockdown (Knockdown) conditions. Blue arcs indicate circRNA isoforms detected under both conditions, whereas red arcs indicate circRNA isoforms detected exclusively following TDP-43 depletion. Arc thickness is proportional to the mean circRNA expression level (CPM) across the four datasets. **B)** RNA-seq quantification of c-circUNC13A expression across the four cellular models under control (CTRL) and TDP-43 knockdown (TDP-43 KD) conditions. Bars represent the mean number of back-splice junction (BSJ)-supporting reads, with individual biological replicates shown as black dots. Error bars indicate ± SD. **C)** RT–qPCR analysis of TDP-43 expression across the four cellular models under control (CTRL) and TDP-43 knockdown (TDP-43 KD) conditions. Transcript abundance was normalised to GAPDH mRNA and is presented as relative expression values (2^−ΔΔCt), with the CTRL condition set to a value of 1. Each dot represents an independent biological replicate (n=4). Statistical significance was assessed using two-tailed unpaired Student's *t* test (***p < 0.001). **D)** RT–qPCR analysis of the indicated transcripts in TDP-43-depleted SK-N-BE(2) cells before (RNase R−) and after (RNase R+) RNase R treatment. Transcript abundance was normalised to a DNA spike-in and is presented as relative expression values (2^−ΔΔCt), with the RNase R-condition set to a value of 1. Each dot represents an independent biological replicate (n=3). Statistical significance was assessed using a two-tailed unpaired Student's *t* test (*p < 0.05,***p < 0.001). **E)** RT–qPCR analysis of the indicated targets in SH-SY5Y cells with a wild-type (Wt) or UNC13A cryptic-exon deleted (ΔCE) background, under control (CTRL) or TDP-43 knockdown (KD) conditions. Transcript abundance was normalised to GAPDH mRNA and is presented as relative expression values (2^−ΔΔCt), with respect to the CTRL samples set to a value of 1. Each dot represents an individual sample (n=4). Statistical significance was assessed using a two-tailed unpaired Student's *t* test (n.s. p > 0.05, **p < 0.01,***p < 0.001).

A

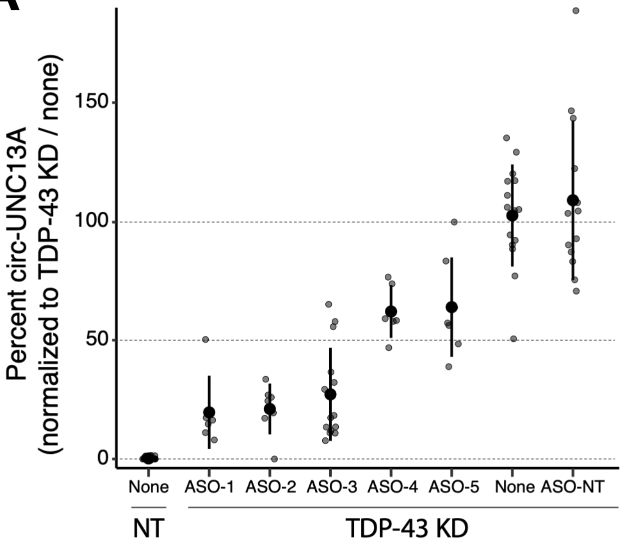
Percent circ-UNC13A
(normalized to TDP-43 KD / none)
150
100
50
0
None
ASO-1
ASO-2
ASO-3
ASO-4
ASO-5
None
ASO-NT
NT
TDP-43 KD

**Figure S12: A)** Percent c-circUNC13A (normalized to TDP-43 knockdown only cells) across different ASO conditions. c-circUNC13A is undetectable in cells without TDP-43 knockdown (NT/none), whereas TDP-43 KD induces c-circUNC13A expression (TDP-43 KD/none). Treatment with splice correcting ASOs (8μM) targeting intron 20 of UNC13A reduces c-circUNC13A levels to varying extents, while a non-targeting control ASO (ASO-NT) has no effect. Individual data points are shown with mean ± SD.
